# Discovery and Characterization of a Pan-betacoronavirus S2-binding antibody

**DOI:** 10.1101/2024.01.15.575741

**Authors:** Nicole V. Johnson, Steven C. Wall, Kevin J. Kramer, Clinton M. Holt, Sivakumar Periasamy, Simone Richardson, Naveenchandra Suryadevara, Emanuele Andreano, Ida Paciello, Giulio Pierleoni, Giulia Piccini, Ying Huang, Pan Ge, James D. Allen, Naoko Uno, Andrea R. Shiakolas, Kelsey A. Pilewski, Rachel S. Nargi, Rachel E. Sutton, Alexandria A. Abu-Shmais, Robert Parks, Barton F. Haynes, Robert H. Carnahan, James E. Crowe, Emanuele Montomoli, Rino Rappuoli, Alexander Bukreyev, Ted M. Ross, Giuseppe A. Sautto, Jason S. McLellan, Ivelin S. Georgiev

## Abstract

Three coronaviruses have spilled over from animal reservoirs into the human population and caused deadly epidemics or pandemics. The continued emergence of coronaviruses highlights the need for pan-coronavirus interventions for effective pandemic preparedness. Here, using LIBRA-seq, we report a panel of 50 coronavirus antibodies isolated from human B cells. Of these antibodies, 54043-5 was shown to bind the S2 subunit of spike proteins from alpha-, beta-, and deltacoronaviruses. A cryo-EM structure of 54043-5 bound to the pre-fusion S2 subunit of the SARS-CoV-2 spike defined an epitope at the apex of S2 that is highly conserved among betacoronaviruses. Although non-neutralizing, 54043-5 induced Fc-dependent antiviral responses, including ADCC and ADCP. In murine SARS-CoV-2 challenge studies, protection against disease was observed after introduction of Leu234Ala, Leu235Ala, and Pro329Gly (LALA-PG) substitutions in the Fc region of 54043-5. Together, these data provide new insights into the protective mechanisms of non-neutralizing antibodies and define a broadly conserved epitope within the S2 subunit.

## INTRODUCTION

Coronaviruses (CoVs) are a broad group of enveloped, positive-sense RNA viruses that can infect a broad spectrum of animals, including pigs, camels, birds, bats, and humans (Li 2016). These viruses have high rates of mutation and recombination frequency that allow for efficient adaptation to a range of hosts (Su, Wong et al. 2016, Amoutzias, Nikolaidis et al. 2022). There are seven known human coronaviruses (HCoVs), of which five belong to the betacoronavirus genus and two belong to the alphacoronavirus genus. Four HCoVs (HCoV-OC43, -229E, - HKU1, and -NL63) cause seasonal respiratory illness with generally mild symptoms that can be more severe in children, the immunocompromised, and the elderly (Nickbakhsh, Ho et al. 2020). Since 2002, three novel betacoronaviruses—severe acute respiratory syndrome (SARS)-CoV, Middle East respiratory syndrome (MERS)-CoV, and SARS-CoV-2—have emerged from animal reservoirs and caused severe disease outbreaks in humans (Ksiazek, Erdman et al. 2003, Rota, Oberste et al. 2003, Zaki, van Boheemen et al. 2012, Zhou, Yang et al. 2020). Most recently, the emergence of SARS-CoV-2 in December 2019 resulted in a global pandemic that has led to nearly 800 million cases and 7 million deaths worldwide to date. These zoonotic spillover events and their devastating effects on the global healthcare system underscore the need for effective countermeasures to future coronavirus outbreaks. Since its emergence, SARS-CoV-2 has continued to circulate, resulting in variants with mutations that enable escape from virtually all clinically approved neutralizing antibodies and decreased vaccine efficacy, emphasizing the urgent need for interventions that provide broad protection from diverse SARS-CoV-2 variants (Hacisuleyman, Hale et al. 2021, Cox, Peacock et al. 2023).

The primary target for coronavirus-neutralizing antibodies is the trimeric spike (S) glycoprotein, which decorates the virion surface and mediates host cell attachment and membrane fusion (Li 2016). Coronavirus spike is a class I fusion protein, expressed as a single polypeptide precursor that requires activation by host cell proteases (Bosch, van der Zee et al. 2003). Proteolysis creates two subunits: an S1 subunit responsible for host cell attachment and tropism, and an S2 subunit that drives membrane fusion. S1 is composed of an N-terminal domain (NTD) and a C-terminal receptor binding domain (RBD) that recognizes a protein receptor on the target cell. The S2 subunit initially folds into a metastable, spring-loaded conformation that contains a globular head composed of the fusion peptide (FP), heptad repeat 1 (HR1), central helix (CH), and connector domain (CD). This region is anchored to the viral membrane by a flexible, elongated helical stalk (Ke, Oton et al. 2020). After cellular attachment, S1 is shed and S2 undergoes conformational rearrangements that drive fusion of the viral and host cell membranes.

The vast majority of neutralizing monoclonal antibodies characterized for SARS-CoV-2 target S1, primarily through epitopes within the RBD (Raybould, Kovaltsuk et al. 2021). S2-directed antibodies are less well characterized, although they comprise a substantial portion of the immune repertoires of convalescent SARS-CoV-2 patients and vaccinated individuals (Amanat, Thapa et al. 2021, Sakharkar, Rappazzo et al. 2021, Voss, Hou et al. 2021). Unlike S1-directed antibodies, S2-directed antibodies tend to be poorly neutralizing, but have increased breadth given the higher conservation of S2 relative to S1. (Grobben, van der Straten et al. 2021, Shiakolas, Kramer et al. 2021, Tong, Gautam et al. 2021, Claireaux, Caniels et al. 2022).

Notably, antibodies that bind SARS-CoV-2 S – mainly within S2 – have been detected in unexposed individuals (Ng, Faulkner et al. 2020, Grobben, van der Straten et al. 2021, Song, He et al. 2021), and some cross-react with S from multiple human coronaviruses and are boosted after SARS-CoV-2 infection or vaccination (Ng, Faulkner et al. 2020, Grobben, van der Straten et al. 2021, Song, He et al. 2021). This suggests that such antibodies were elicited by one of the four circulating human coronaviruses, indicating that S2 may be an important immunogen for pan-coronavirus vaccine efforts.

To date, three major groups of broadly reactive S2-directed antibodies have been characterized. The largest group – isolated from human donors or vaccinated animals – binds to the stem helix, just C-terminal to the globular head, and tend to be weakly or non-neutralizing with varying potency across multiple coronaviruses (Hsieh, Werner et al. 2021, Sauer, Tortorici et al. 2021, Wang, van Haperen et al. 2021). However, many broadly reactive, stem helix-directed antibodies offer some degree of protection against SARS-CoV-2 challenge in mice and Syrian hamsters (Hsieh, Werner et al. 2021, Pinto, Sauer et al. 2021, Zhou, Yuan et al. 2022, Dacon, Peng et al. 2023, Zhou, Song et al. 2023). The second group of S2-directed antibodies targets the fusion peptide (Dacon, Tucker et al. 2022, Low, Jerak et al. 2022, Sun, Yi et al. 2022). Like the stem-helix binders, antibodies that target the fusion peptide are generally weakly neutralizing but offer prophylactic protection against SARS-CoV-2 in mice and hamsters (Dacon, Tucker et al. 2022, Low, Jerak et al. 2022). A third group of S2-directed antibodies targets the membrane-distal apex of S2, a region that is likely exposed through transient trimer opening or after S1 shedding (Chen, Gilchuk et al. 2021, Claireaux, Caniels et al. 2022, Costello, Shoemaker et al. 2022, Silva, Huang et al. 2023). None of these apex-directed antibodies have been shown to neutralize authentic SARS-CoV-2 virus, and their protective capacity in animal models has not been determined (Claireaux, Caniels et al. 2022, Silva, Huang et al. 2023).

Here, we used LInking B cell Receptor to Antigen specificity through sequencing (LIBRA-seq) to examine the B cell repertoire of a convalescent COVID-19 donor for broadly reactive coronavirus spike-directed antibodies. We identified 54043-5, a non-neutralizing antibody that binds to a large panel of betacoronavirus spike proteins, including those that infect humans. 54043-5 utilizes an uncommon gene pairing and targets a cryptic epitope at the apex of the S2 subunit. Although we found the wild-type IgG to be non-protective in murine models, addition of an Fc-silencing mutation (LALA-PG) led to some protection in these mice. The characterization of 54043-5 offers insight into the strategic targeting of the S2 subunit for broad interventions to betacoronavirus infection and the role that non-neutralizing antibodies play in protection from disease.

## RESULTS

### Identification and characterization of broadly reactive coronavirus antibodies by LIBRA-seq

To identify B cells that are broadly reactive to diverse coronavirus spike proteins, we utilized the LIBRA-seq technology for antibody discovery (Setliff, Shiakolas et al. 2019). Using this method, a LIBRA-seq score is calculated for each B cell receptor (BCR) based on antigen binding that estimates the strength of the paired interaction. We performed LIBRA-seq on samples from two adult donors, one of whom had previously recovered from a SARS-CoV-2 infection. We previously described the isolation and characterization of a SARS-CoV-2 spike-directed antibody, 54042-4, from the same experiments (Kramer, Johnson et al. 2021). Here, we mined the LIBRA-seq data for B cells that exhibited high breadth of reactivity against diverse coronavirus spikes (**Figure 1A**). Several B cells exhibited high LIBRA-seq scores for multiple probes, indicative of broad spike reactivity, particularly to spikes from SARS-CoV-2, SARS-CoV, and MERS-CoV (**Figure 1B**). Based on these results, the BCR sequences for 50 predicted cross-reactive B cells were selected for validation as monoclonal antibodies, which were cloned, expressed in microscale, purified, and screened for binding (**Figure S1A,B**). Spike cross-reactivity was confirmed for several antibodies that were associated with diverse sequence features, including diverse heavy and light chain V-genes, and complementarity-determining region 3 (CDR3) length and composition, underlining a wide range of possible mechanisms for spike cross-reactivity (**Figure 1C**). As expected based on spike sequence similarity, the majority of cross-reactive antibodies bound to spikes from SARS-CoV-2 and SARS-CoV, although a subset additionally bound to spikes from HCoV-OC43 and HCoV-HKU1 (antibodies 54042-13 and 54043-5) and/or MERS-CoV (54041-1, 54043-4, and 54043-5) (**Figure 1C)**. Among these, antibody 54043-5 exhibited the greatest breadth and was selected for further characterization.

**Figure 1.**
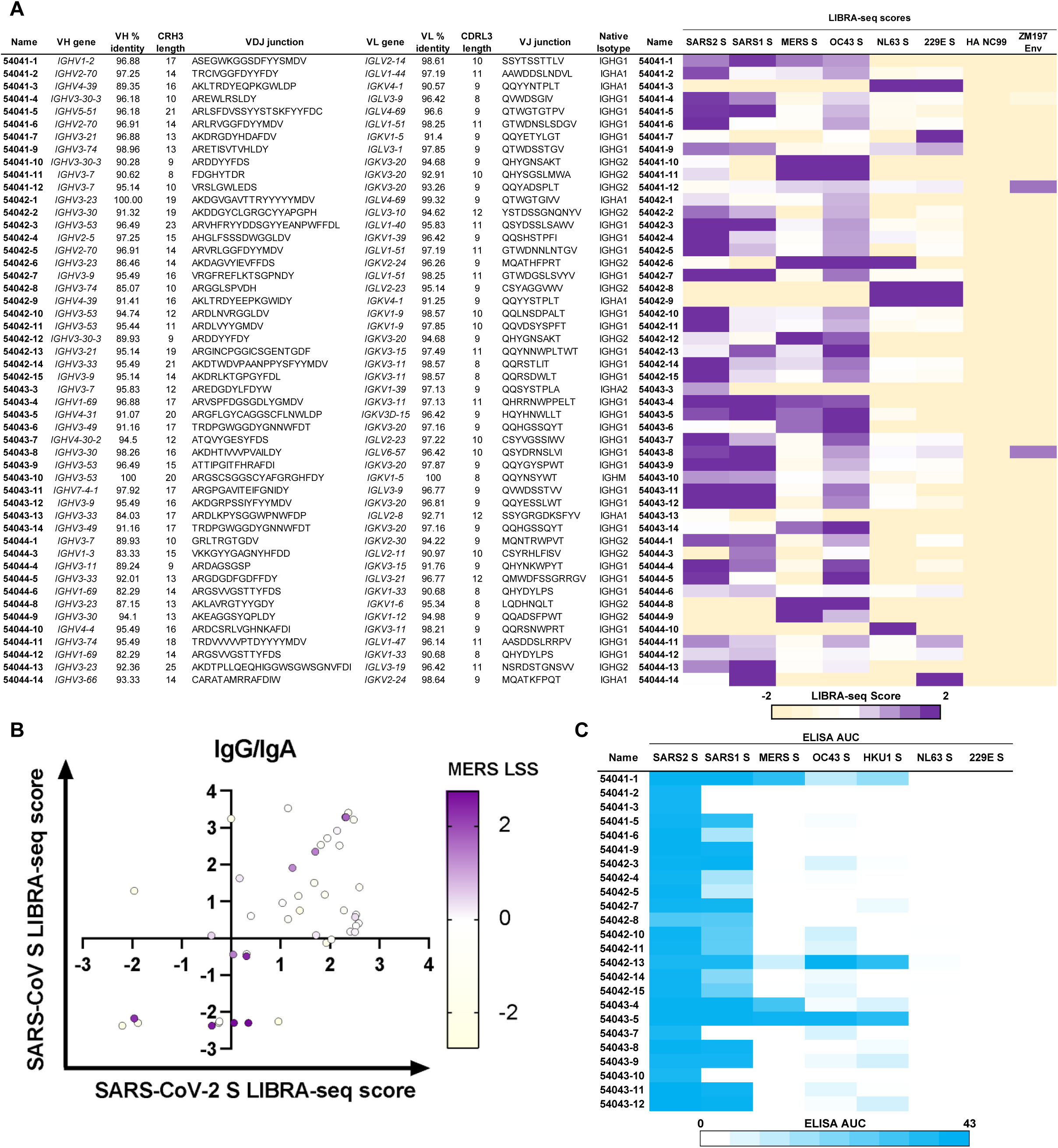
Identifying broadly reactive mAbs using a diverse CoV spike LIBRA-seq panel. **(A)** A panel of antibodies identified by LIBRA-seq (rows), with corresponding V- and J-genes and percent nucleotide identities, CDR lengths and amino acid sequences, and isotype (columns). LIBRA-seq scores for each antigen are shown alongside each antibody as a heatmap from -2 (tan) to 2 (purple). HA-NC99 was included as a negative control antigen. **(B)** 50 IgG and IgA cells identified by LIBRA-seq are shown as circles, with their respective LIBRA-seq scores (LSSs) for SARS-CoV-2 spike (x axis), SARS-CoV spike (y axis), and MERS-CoV spike (color heatmap). **(C)** Monoclonal antibodies were produced by microexpression and tested for binding by ELISA to a panel of human coronavirus spike proteins. ELISA area under the curve (AUC) values were calculated from binding curves in Figure S1A and are shown as heatmaps from minimum (white) to maximum (blue) binding. Antibodies that bound to the SARS-CoV-2 spike are shown. ELISA controls are described in Figure S1.

### Antibody 54043-5 is broadly reactive and targets the S2 subunit

To further assess the breadth of reactivity for antibody 54043-5, we tested its binding to an extended panel of purified spike proteins from several coronavirus genera (**Figure 2A**). As before, antibody 54043-5 displayed high reactivity for all the human betacoronavirus spike antigens tested (SARS-CoV, SARS-CoV-2, MERS-CoV, HCoV-OC43, and HCoV-HKU1), with ELISA area under the curve (AUC) values between 39.3 and 41.1, whereas no binding was observed for the human alphacoronavirus spikes from HCoV-NL63 and HCoV-229E (**Figure 2B**). We also tested binding to five non-human coronavirus spike proteins including two from bat betacoronaviruses and three from porcine coronaviruses from other genera. The WIV1-CoV spike, from a bat betacoronavirus belonging to the same lineage as SARS-CoV-2, was bound by 54043-5 at a similar half maximal effective concentration (EC50) to that of SARS-CoV-2 (3.2 ng/mL and 3.3 ng/mL respectively). The other bat betacoronavirus, HKU9-CoV, is from the distant D lineage of the betacoronavirus genus and was the only betacoronavirus tested whose spike showed negligible binding by 54043-5 at the highest concentration tested (10 μg/mL).

**Figure 2.**
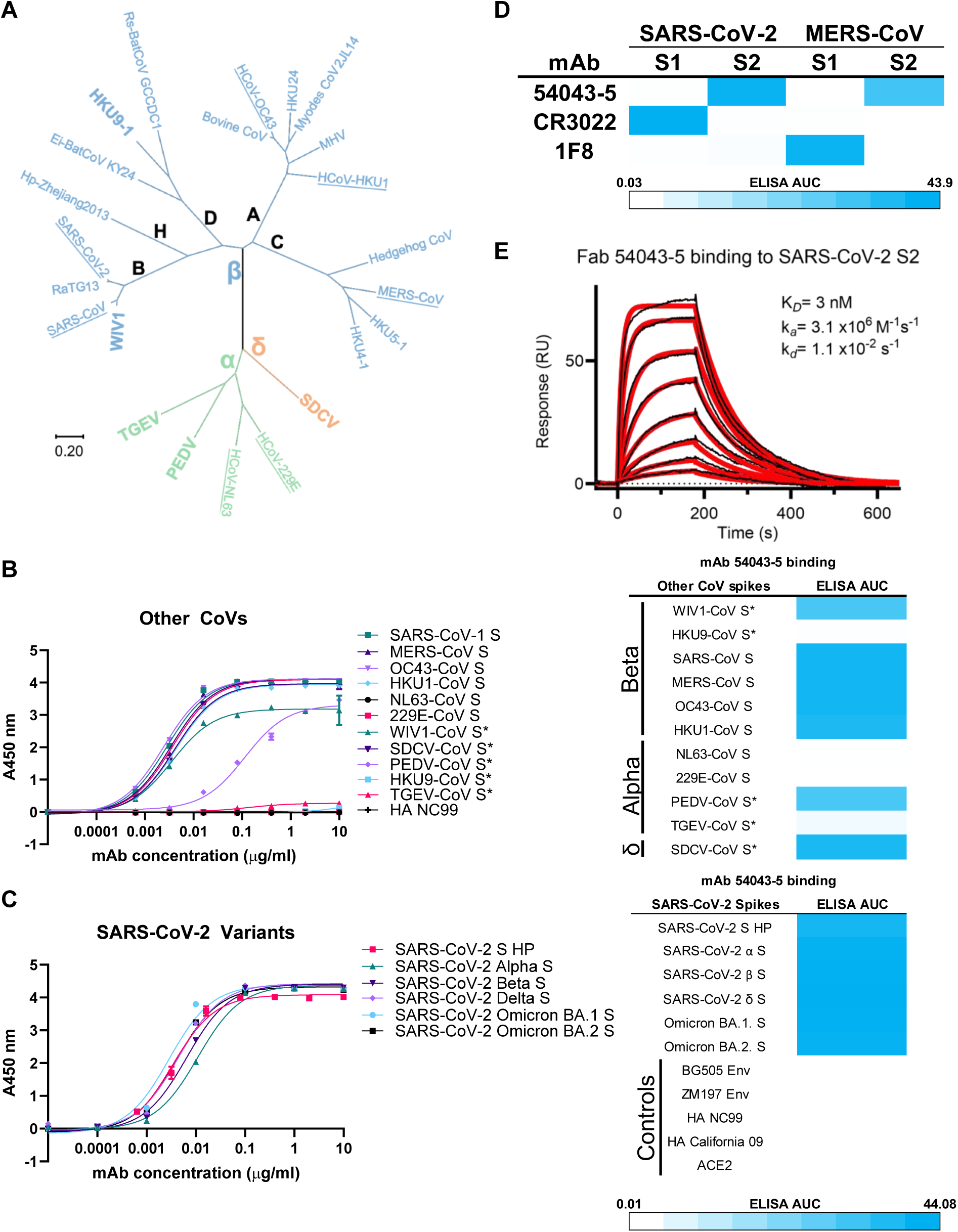
mAb 54043-5 is an ultrabroad S2 antibody. **(A)** Phylogenetic tree of coronavirus spikes from select members of the *Orthocoronavirinae* subfamily. Members of the alpha- (α) beta- (β) and deltacoronavirus (δ) genera are indicated by color. The betacoronavirus genus contains five lineages: A-D and the Hibecovirus lineage (H). Spikes included in LIBRA-seq experiments and binding assays are underlined, and those included exclusively in binding assays are boldly labeled. Scale bar denotes amino acid phylogenetic distance. **(B)** ELISA curves for mAb 54043-5 binding to spikes from selected CoVs (left) and associated ELISA AUC values for each (right). An “*” denotes a non-human coronavirus **(C)** ELISA curves for mAb 54043-5 binding to spikes from SARS-CoV-2 variants (left) and associated ELISA AUC values for each (right) **(D)** ELISA AUCs for 54043-5 binding to the S1 and S2 domains of spikes from SARS-CoV-2 and MERS-CoV, or positive control mAbs CR3022 and 1F8. ELISA controls described in figure S1 **(E)** SPR sensorgram for 54043-5 Fab binding to the S2 subunit of the SARS-CoV-2 spike. Binding curves are colored black, and data fit to a 1:1 binding model is colored red.

Interestingly, the HKU9-CoV spike shares 31.4% amino acid identity with the SARS-CoV-2 spike, which is higher than three of the human betacoronavirus spikes that 54043-5 binds, namely those from MERS-CoV (31.2%), HCoV-OC43 (30.7%), and HCoV-HKU1 (29.8 %) (**Table S1**). Spikes from the porcine TGEV-CoV, PEDV-CoV, and SDCV-CoV showed variable reactivity to 54043-5. Among the two alphacoronaviruses, TGEV-CoV spike displayed negligible binding at high concentrations of 54043-5 antibody, whereas the PEDV-CoV spike showed robust, albeit reduced, binding with an ELISA AUC of 31.5. The deltacoronavirus SDCV-CoV spike reactivity closely resembled that of the human betacoronaviruses despite sharing only 27.4% amino acid identity with SARS-CoV-2 spike, with an ELISA AUC of 39.5. Together, we determined 54043-5 to exhibit a high breadth of reactivity to a diverse set of spike proteins from human and non-human betacoronaviruses, with additional reactivity to some alpha- and deltacoronavirus spikes. Furthermore, the percent sequence identity shared with the SARS-CoV-2 spike does not appear to be predictive of 54043-5 reactivity among the spikes tested.

As SARS-CoV-2 variants of concern continue to emerge, we next assessed the reactivity of antibody 54043-5 to five SARS-CoV-2 variants: Alpha, Beta, and Delta, as well as Omicron BA.1 and BA.2 by ELISA (**Figure 2C**). The results revealed robust binding to spikes derived from all five SARS-CoV-2 variants tested, suggesting that 54043-5 binding to SARS-CoV-2 spikes is unaffected by substitutions within the spike antigens from these variants of concern.

To narrow down the region of the spike targeted by 54043-5, we performed ELISA binding assays using SARS-CoV-2 and MERS-CoV spike antigens that contain only the S1 or S2 subunit (**Figures 2D, S1C**). Antibodies CR3022 (ter Meulen, van den Brink et al. 2006, Yuan, Wu et al. 2020) and 1F8 (Tang, Agnihothram et al. 2014) were used as S1-binding controls for SARS-CoV-2 and MERS-CoV, respectively. As expected, we observed binding of the control antibodies to S1, but not S2 of their target spike proteins. In contrast, 54043-5 bound to S2, but not S1, of both SARS-CoV-2 and MERS-CoV spikes. To determine the binding affinity of 54043-5 to SARS-CoV-2 S2, surface plasmon resonance experiments were performed by immobilizing a prefusion-stabilized S2 protein (Hsieh, Zhou et al., in preparation) to a Ni-NTA chip via its C-terminal His tag and flowing over various concentrations of the 54043-5 antigen-binding fragment (Fab). The resulting association and dissociation curves were fit to a 1:1 binding model, resulting in a calculated *K*_D_ of 3 nM (**Figure 2E**). Collectively, these results demonstrate that 54043-5 is a high-affinity, S2-directed antibody.

Since in some cases antibody cross-reactivity with diverse antigens may be due to promiscuous polyreactivity, we next evaluated the reactivity of antibody 54043-5 against a well-established panel of auto-antigens (Yang, Holl et al. 2013) (**Figure S1D**). These experiments also included two additional LIBRA-seq antibodies with high breadth of coronavirus spike reactivity, 54041-1 and 54043-4 (**Figure 1C**). The three antibodies showed negligible reactivity to all of the auto-antigens and other non-coronavirus antigens (**Figure S1B,D**), indicating that the breadth of antigen reactivity for these antibodies is due to specific binding to coronavirus spike proteins.

### 54043-5 binds the apex of S2

To elucidate the epitope of 54043-5, we solved a 3.0 Å resolution structure of the 54043-5 Fab bound to the SARS-CoV-2 S2 subunit by cryo-EM (Hsieh, Zhou et al., in preparation)(**Figures 3A, S2; Table S2**). Classification of extracted particles yielded 2D class averages exhibiting top and side views of the S2 subunit with three Fabs bound. 3D reconstruction yielded a single volume for the complex, representing S2 in a closed state with the bound Fabs tightly packed at the apex of the trimer. The refined structure revealed an epitope within a single protomer that spans the helix-turn-helix formed at the junction of heptad repeat 1 (HR1) and the central helix (CH). 54043-5 buries a total surface area of 629 Å^2^ on S2 at an interface dominated by the CDR3s of the heavy and light chains, which grip the region between S2 residues Asn969 and Arg995 (**Figures 3B, S3A-C; Table S3)**.

**Figure 3.**
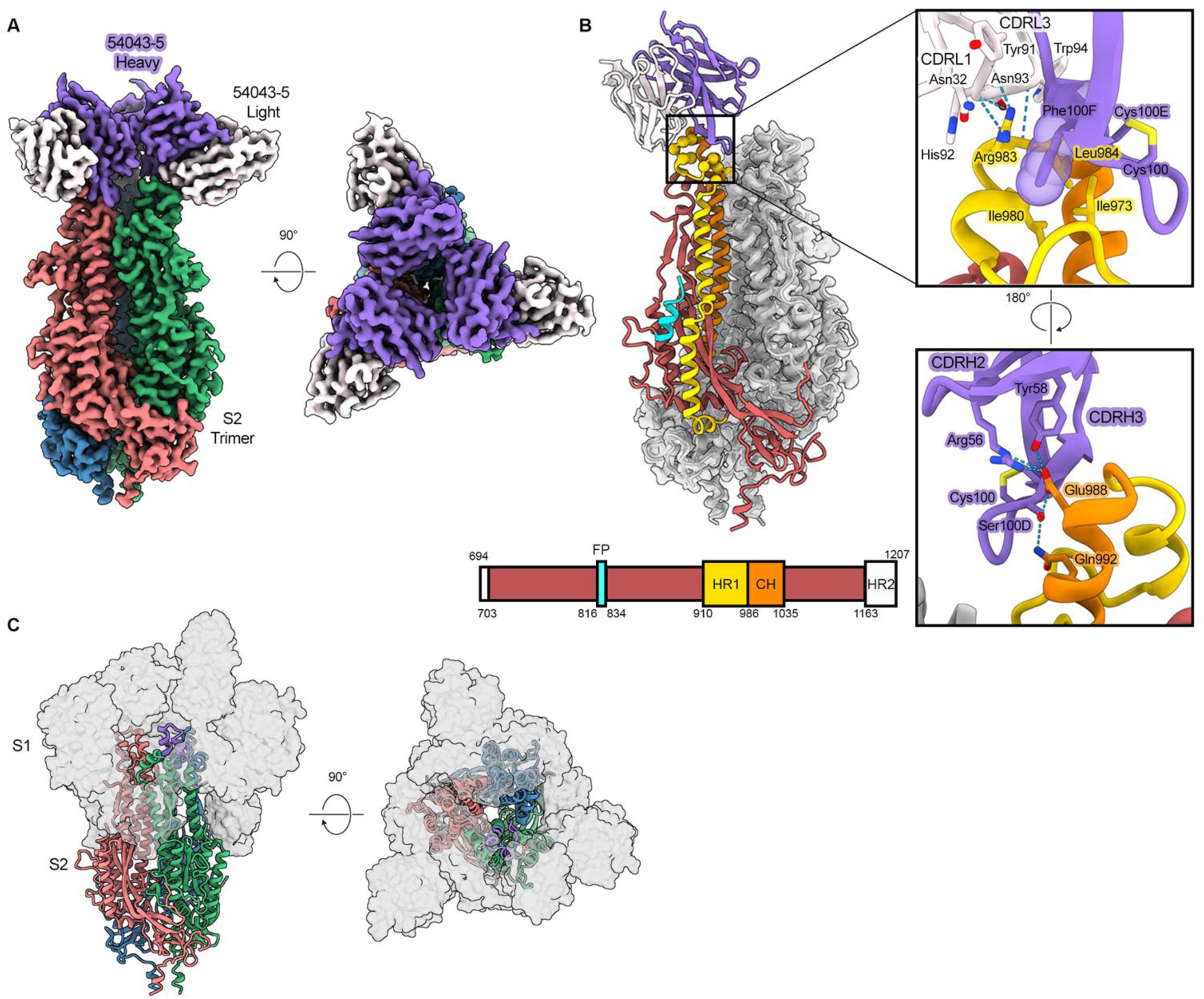
Cryo-EM structure of Fab 54043-5 bound to the SARS-CoV-2 S2 subunit. **(A)** Side and top-down views of the 3.0 Å 3D reconstruction of Fab 54043-5 bound to S2. The S2 protomers are colored green, blue, or salmon. The 54043-5 heavy and light chains are colored purple and white, respectively. **(B)** 54043-5 binds an epitope at the apex of S2, spanning the junction between heptad repeat 1 (HR1) and the central helix (CH). The S2 subunit bound to one Fab is shown as cartoons (left), with two protomers underneath the corresponding EM map and colored gray. One protomer is colored according to the gene schematic (bottom) and 54043-5 is colored as in (A). Zoomed in views of the 54043-5 Fab-S2 interface (right) are shown as cartoons, with important residues shown as sticks. Phe100F is shown with a partially transparent surface to illustrate space filling in a hydrophobic pocket within the epitope. Oxygen atoms are colored red, nitrogen atoms are colored blue, sulfur atoms are colored yellow, and hydrogen bonds are shown as light blue dashed lines. **(C)** Top and side views of the SARS-CoV-2 spike protein, with the S2 subunit shown as cartoons and colored as in (A). The 54043-5 epitope on one protomer is colored in purple. The S1 subunit, shown as a partially transparent gray surface, almost entirely covers the apex of S2 and the 54043-5 epitope when present.

The heavy chain buries 402 Å^2^ on the S2 protomer primarily through the extended, largely nonpolar CDRH3 loop, which is stabilized by a disulfide bond between Cys100 and Cys100E (Kabat numbering). The CDRH3 reaches toward the center of the trimer, contacting nearly every residue within the epitope. Of note, Phe100F is buried in a hydrophobic pocket formed by Ile973, Ile980, and Leu984 within HR1 and capped by a cation-pi interaction with the sidechain of Arg983. Proximal to the trimeric interface, Ser100D forms a sidechain hydrogen bond with CH residues Glu988 and Gln992. CDRH2 contributes to the interface just two residues, Arg56 and Tyr58, which represent two somatic mutations within the VH4-31 gene. Each form additional polar interactions with Glu988, including two possible salt bridges with Arg56 and a hydrogen bond with Tyr58.

The 54043-5 light chain contributes 227 Å^2^ to the interface through contacts at the junction between HR1 and CH including residues Ser982 through Asp985. The light chain utilizes only five residues for this interaction: four from CDRL3 and one from CDRL1. A striking number of contacts involving each of these residues occur with Arg983 of HR1. Here, a backbone hydrogen bond is formed with CDRL3 residue Trp94, and the Arg983 sidechain forms three additional hydrogen bonds with the backbone atoms of Tyr91 and His92. Further, several contacts with Arg983 are mediated by the sidechains of Tyr91, His92, Asn93 and CDRL1 residue Asn32, which join CDRH3 residue Phe100F to fully surround the residue.

Interestingly, the S2 epitope bound by 54043-5 is inaccessible in the closed, prefusion spike. Prior to S1 shedding, the S1 subunit caps S2, surrounding the HR1 and CH helices (**Figure 3C**). Although our LIBRA-seq and ELISA data show that 54043-5 binds to a prefusion-stabilized SARS-CoV-2 spike (S-6P), we were unable to obtain a structure of the complex. This suggests that binding does not occur to the intact, closed prefusion conformation, but rather during transient ‘breathing’ of the spike trimer that has been shown to allow for binding of murine antibody 3A3, which binds a similar epitope (**Figure S2D**) (Costello, Shoemaker et al. 2022, Silva, Huang et al. 2023).

### 54043-5 has uncommon genetic features and binds an epitope that is highly conserved across betacoronaviruses

The discovery that 54043-5 targets a cryptic epitope and broadly binds coronavirus spikes led us to investigate its gene pairing and CDR sequence features within the broader context of spike-directed antibodies in human repertoires. We searched the CoV-AbDab database, which aggregates paired heavy- and light-chain sequences of coronavirus antibodies and found that only a small fraction (6 out of 9,829 as of February 21, 2023) share the same VH (*IGHV4-31*) and VL (*IGKV3D-15*) genes as 54043-5 (Raybould, Kovaltsuk et al. 2021). Among these, only two antibodies showed a high sequence identity (>50%) to the CDRL3 and zero showed a high sequence identity to the CDRH3 of 54043-5 (**Figure 4A**). In contrast, other apical S2-directed antibodies identified were part of a common public clonotype utilizing the *IGHV1-69*/*IGKV3-11* gene pairing (Claireaux, Caniels et al. 2022).

**Figure 4.**
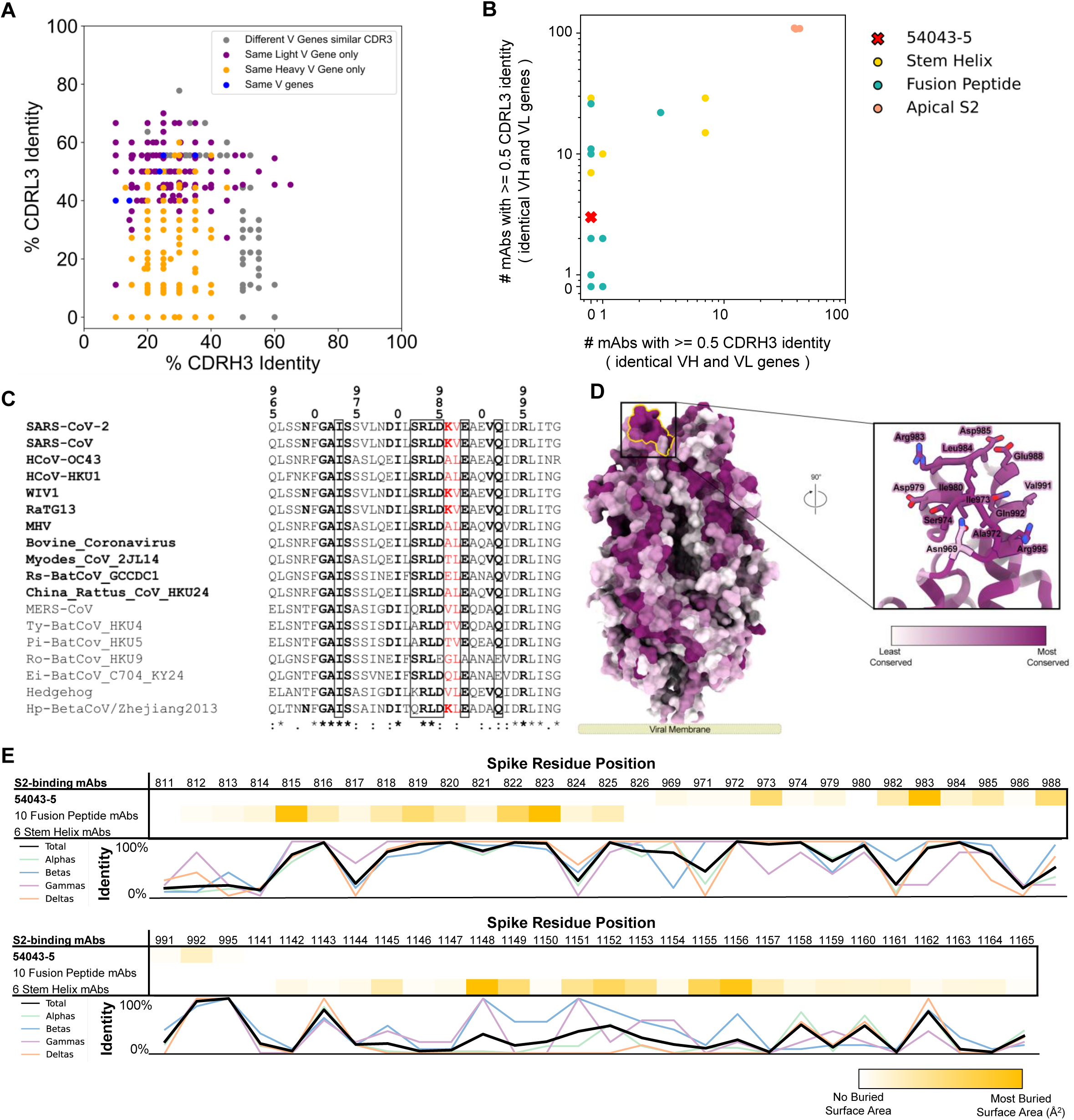
Antibody sequence feature uniqueness and conservation of epitope across CoVs. **(A)** Sequence feature analysis of mAb 54043-5 compared to coronavirus antibodies in the CoVAbDab. The plot displays the percent amino acid identity of the CDRH3 (x-axis) and CDRL3 (y-axis) of a subset of antibodies to 54043-5. Antibodies shown have at least one identical variable (V) gene or ≥50% CDR3 sequence identity. Colors denote shared V gene usage with 54043-5. **(B)** The count of similar public clones for each S2-binding antibody with characterized epitopes, based on epitope group. Each antibody is represented as a point, with its x-axis coordinate reflecting the number of antibodies in the CoVAbDab sharing both heavy and light V genes and having a CDRH3 with ≥50% amino acid sequence identity. The y-axis coordinate corresponds to the number of antibodies with the same V genes and a CDRL3 sequence with ≥50% identity. **(C)** Sequence alignment of Betacoronavirus strains, focused on the epitope bound by mAb 54043-5. Structurally buried residues are in bold, with those significantly buried (≥25 Å^2^ buried solvent accessible surface area) enclosed in boxes. S2P mutation residues are highlighted in red, and strains with complete conservation of significantly buried residues are listed in bold. **(D)** Sequence conservation within the S2 subunit of the spike, mapped onto a surface representation of the S2 subunit and colored based on sequence alignment in (C) (left). The 54043-5 epitope is outlined in yellow. A zoomed in view of the S2 apex is shown as cartoons, with 54043-5 epitope residues shown as sticks. **(E)** Relative epitope buried surface area (BSA) and conservation across the Orthocoronavirinae genera. Two conserved and structurally characterized S2-epitopes are shown compared to mAb 54043-5, with relative BSA shown as a color gradient from white (0 BSA) to dark green (most BSA within the row). The percent conservation of each epitope residue among the four genera are shown as a line graph.

To put these results into context, we analyzed these relationships within the two other structurally characterized groups of S2-directed antibodies (**Figures 4B, S4)**. Within the first group, composed of stem-helix-directed antibodies, we found that each antibody queried had dozens of other antibodies within the CoV-AbDab database with similar CDRL3s (>50% sequence identity) and identical VH/VL genes, but only two stem-helix antibodies had a large number of matches with similar CDRH3s and identical VH/VL genes. Fusion peptide-directed antibodies displayed the rarest sequence features among the three groups, though two of these antibodies (DH1058 and COV91-27) had at least 20 counterparts in the database sharing the same VH/VL genes and similar CDRL3s. Compared to these groups, apical S2-antibodies were the most common, but 54043-5 stood out from other characterized antibodies within this group by having rare sequence features comparable to levels observed for fusion peptide antibodies.

To visualize the extent to which the 54043-5 epitope is conserved across the betacoronavirus genus, we compiled an alignment of betacoronavirus spike protein sequences, focusing on the residues within the 54043-5 epitope. (**Figure 4C**). Only one species, *Rousettus bat coronavirus HKU9*, contained more than one substitution in the significantly buried epitope residues (BSA ≥25 Å^2^), providing a likely explanation for the lack of binding observed between the HKU9 spike and 54043-5 (**Figure 2B**). Mapping the sequence conservation onto the structure of SARS-CoV-2 S2, using a color gradient to indicate high (purple) or low (white) levels of conservation, further revealed that the 54043-5 epitope contains some of the most highly conserved residues within the S2 subunit (**Figure 4D**). The only exception is residue Asn969, which lies on the periphery of the epitope and is not significantly buried by 54043-5. The structural conservation analysis also juxtaposes residues included in the 54043-5 epitope with excluded residues interspersed along the HR1/CH junction, which largely exhibit much lower levels of conservation.

We next compared the overall conservation of the 54043-5 epitope to the epitopes of other structurally characterized S2 antibodies. Spike residues within the epitopes of 54043-5, 10 fusion peptide-, or 6 stem-helix-directed antibodies were defined based on relative buried surface area and analyzed for conservation across all genera within the *Orthocoronavirinae* subfamily (**Figure 4E**). Five of the seven 54043-5 epitope residues with significantly buried surface area— 973, 983, 984, 985, and 992—showed high conservation among all CoVs. The remaining two residues, 982 and 988, were well conserved among betacoronaviruses, but were generally less conserved in the other genera. Comparatively, epitopes targeted by fusion peptide-directed antibodies exhibited similarly high levels of conservation among all coronaviruses, whereas the epitopes for stem–helix-directed antibodies were generally less well conserved, particularly among coronaviruses outside the betacoronavirus genus. Together, these results indicate that antibody 54043-5 utilizes unique sequence features to target the highly conserved apical S2 epitope, which is more commonly targeted than the similarly conserved fusion peptide epitope and less conserved stem-helix epitope.

### 54043-5 is a non-neutralizing antibody that induces Fc effector functions

We tested antibody 54043-5 for neutralization of pseudotyped or authentic SARS-CoV-2 and found it to be non-neutralizing in both experiments (**Figure S5**). Because non-neutralizing antibodies may offer protection through the induction of Fc-dependent antiviral activity, we next tested 54043-5 and additional cross-reactive antibodies from the same individual (**Figure 1C**) to determine if they could induce antibody-dependent cellular phagocytosis (ADCP), antibody-dependent cellular cytotoxicity (ADCC) and antibody-dependent cellular trogocytosis (ADCT). Antibodies 54041-1, 54043-4 and 54043-5, but not 54042-13, effectively triggered the uptake of beads coated with SARS-CoV-2 D614G spike by THP-1 cells (**Figures 5A, S5E**). In a similar assay testing phagocytosis by primary human monocytes (ADMP), only 54043-4 triggered phagocytosis to a greater extent than the negative control with no antibody included, whereas 54041-1 and 54043-5 (54042-13 was not tested) showed very low levels of ADMP activity (**Figure 5B**). We next tested ADCT by incubating HEK293T “donor” cells expressing biotinylated SARS-CoV-2 spikes with stained THP-1 “recipient” cells. Trogocytosis was measured by flow cytometry using Streptavidin-PE to detect the transfer of biotinylated spikes to the surface of THP-1 cells. Similar to what we observed with ADCP, 54041-1, 54043-4 and 54043-5, but not 54042-13, induced high levels of trogocytosis (**Figures 5C, S4G**). When incubated with neutrophils, only 54041-1 induced a cytotoxic response (**Figures 5D, S4F**). We also tested whether these cross-reactive antibodies could induce ADCP of beads coated with HCoV-OC43 spike and found that all four triggered high levels of phagocytosis, though 54041-1 did so to a lesser extent (**Figures 5E, S4H**). Together, these data demonstrate a diverse range of Fc effector phenotypes for this set of cross-reactive antibodies.

**Figure 5.**
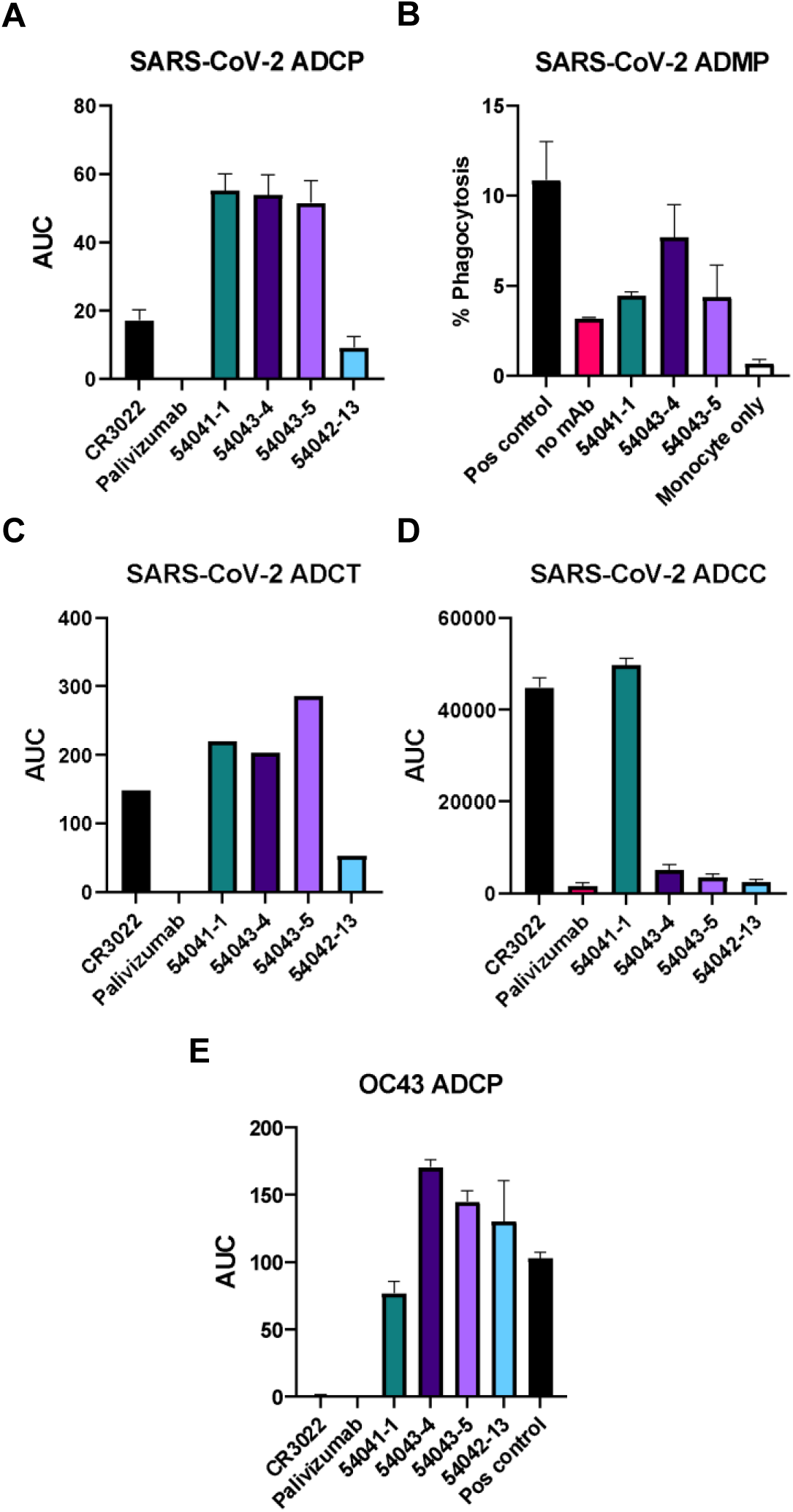
Fc effector functional characteristics of lead candidates in bead-based and cell-based assays. **(A)** Lead cross-reactive mAb candidates, 54041-1, 54043-4, 54043-5, and 54042-13 were tested for their ability to mediate antibody-dependent cellular phagocytosis (ADCP) for SARS-CoV-2, compared to antigen-positive control CR3022 and negative control palivizumab (an anti-RSV antibody). AUC shown was calculated based on the phagocytosis score in figure S4E. **(B)** 54041-1, 54043-4, and 54043-5 were tested for their ability to mediate antibody-dependent monocyte phagocytosis (ADMP) for SARS-CoV-2, compared to a no-antibody negative control. CR3022 was used as an antigen-positive control. Percent phagocytosis calculation is detailed in the methods section. **(C)** Cross-reactive mAbs were tested for their ability to mediate antibody-dependent cellular trogocytosis (ADCT) for SARS-CoV-2, compared to antigen-positive control CR3022 and negative control palivizumab. AUC shown was calculated based on the trogocytosis score in figure S4G. **(D)** Cross-reactive mAbs were tested for their ability to mediate antibody-dependent cellular cytotoxicity (ADCC) against SARS-CoV-2, compared to antigen-positive control CR3022 and negative control palivizumab. AUC shown was calculated based on the cytotoxicity score in figure S4F. **(E)**. Cross-reactive mAbs were additionally tested against OC43 for their ability to mediate antibody-dependent cellular phagocytosis compared to the OC43-specific positive control, 54044-5, and negative control palivizumab. AUC shown was calculated based on the phagocytosis score in figure S4H. All Fc effector data is shown as mean ±SDs.

### 54043-5 LALA-PG administered prophylactically partially protects mice from lethal SARS-CoV-2 challenge

To evaluate the *in vivo* function of 54043-5, we performed a prophylactic study in K18-hACE2 transgenic mice (**Figures 6A-C and S6A**). Twenty-four hours prior to challenge with SARS-CoV-2, mice (n=8 per group) were administered with 12 mg/kg of 54043-5, 54043-5 harboring mutations that have been shown to significantly reduce binding of complement and cell-mediated cytotoxicity (54043-5 LALA-PG), or an IgG1 isotype control antibody (#1664) (Forgacs, Abreu et al. 2021). An additional group was administered 5 mg/kg of neutralizing antibody S309 as a positive control (Pinto, Park et al. 2020). Mice treated with 54043-5 experienced similar weight loss to that of the vehicle and isotype control groups, losing 25% on average of their original weight within 8 days post-infection (p.i.) (**Figure 6A)**. In contrast, the group treated with 54043-5 LALA-PG lost weight for the first 8 days (losing 19% on average of their original weight) and surviving animals recovered to their original weight by day 10. Additionally, the weight loss experienced during days 5–7 was significantly lower (p = 0.0429, 0.0141, 0.0019, respectively) compared to the vehicle administered group (**Figure S6B**). Consistent with the body weight data, none of the mice treated with 54043-5, isotype control antibody, or vehicle control survived past day 10, whereas 40% of mice treated with 54043-5 LALA-PG and 100% of those treated with antibody S309 survived through the end of the study (**Figure 6B**).

**Figure 6.**
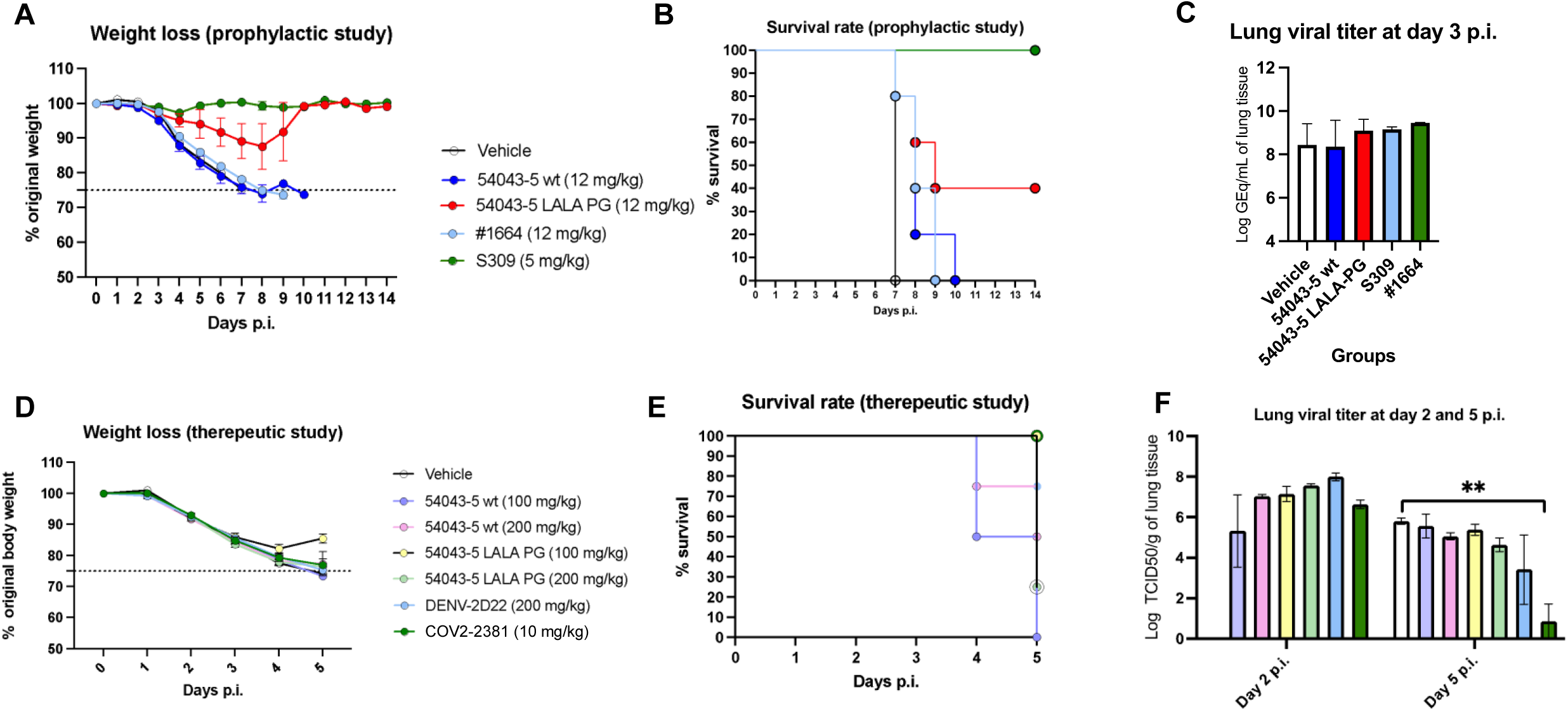
Treatment of mice with 54043-5 as pre- or post-exposure treatment of SARS-CoV-2 infection. Mice were treated, according to group, either 24 hours prior to (prophylactic) or following (therapeutic) intanasal SARS-CoV-2 challenge. Results are expressed as absolute mean values plus SEM. **(A, B)** Body weight **(A)** and survival **(B)** was monitored daily for K18-hACE2 mice treated prophylactically with PBS buffer (vehicle), wild-type 54043-5, 54043-5 LALA-PG, an isotype control (#1664), or a neutralizing SARS-CoV-2 antibody (S309) for 14 days following challenge with SARS-CoV-2 (WA1/2020). Statistical significance of the body weight differences between the mAb testing groups and the vehicle group at days 5-7 p.i. are reported in Figure S6B. **(C)** Three mice per group in the prophylactic study were sacrificed 3 days post-infection (p.i.) and viral titers in the lung tissue were measured. Lung viral titer is expressed as the logarithm of the viral genomic equivalents (GEq) per mL of homogenized lung tissue. **(D, E)** Body weight **(D)** and survival **(E)** were monitored daily for BALB/c mice treated therapeutically with PBS buffer (vehicle), wild-type 54043-5, 540435-LALA-PG, an isotype control antibody (DENV-2D22), or a neutralizing SARS-CoV-2 antibody (COV2-2381) for five days following challenge with SARS-CoV-2 (MA10). **(F)** Four mice per group in the therapeutic study were sacrificed on days 2 and 5 p.i. and viral titers in the lung tissue were measured. Lung viral titer is expressed as the logarithm of the 50% tissue culture infectious dose of virus (TCID50) per gram of lung tissue. **, p<0.01.

Lung samples from a set of mice from the same treatment groups (n=3 per group) were collected at 3 days p.i. to assess their lung viral titers by qRT-PCR. Mice prophylactically administered S309 and 54043-5 LALA-PG antibodies showed average lung viral titers of 1.44×10^9^ and 1.24×10^9^ viral genomic equivalents per milliliter of lung tissue (Geq/mL), respectively (**Figure 6C**). In contrast, mice prophylactically administered the isotype control (#1664) and the 54043-5 antibodies showed a higher average lung viral titer of 2.83×10^9^ and 2.27×10^8^ Geq/mL, respectively. No significant differences between lung viral titers among any groups were observed.

### 54043-5 LALA-PG administered therapeutically partially protect mice from lethal SARS-CoV-2 challenge

We next assessed the protective efficacy of antibody 54043-5 administered therapeutically 24 hours after challenge with SARS-CoV-2 (**Figures 6D-F, S6A**). Groups of BALB/cAnNHsd mice were administered 100 mg/kg of either 54043-5 or 54043-5 LALA-PG, 200 mg/kg of an isotype control antibody (DENV-2D22) (de Alwis, Smith et al. 2012), or 10 mg/kg of SARS-CoV-2 neutralizing antibody (COV2-2381) and were monitored for 5 days. The group treated with 54043-5 LALA-PG experienced the least amount of weight loss during the observation period, corresponding to a total average loss of 12% of their original body weight and a striking 100% survival by day 5 p.i. (**Figure 6D,E**). However, some protection was also observed in mice administered the isotype control antibody DENV-2D22, corresponding to an average weight loss of 20% and an average 75% survival. Conversely, mice treated with 54043-5 showed no significant improvement over the vehicle control, losing on average 25% of their original body weight and completely succumbing to disease by day 5 p.i.

We also assessed the severity of disease in these mice by clinical score, which considers changes in general condition (fur, eyes, posture), motility, and respiration in addition to change in body weight (Mann, Vahle et al. 2012)(**Figure S6C**). After clinical assessment, mice were assigned a score between 0 (no disease burden) to 4 (severe burden, humane endpoint). All groups exhibited an increase in their clinical score through day 3 p.i. Through days 4 and 5, the clinical scores of groups treated with either the vehicle or 54043-5 continued to increase. Similarly, the group treated with DENV-2D22 had an increased clinical score through day 4, with a slight decrease on day 5. Mice treated with COV-2381 reached their maximum mean clinical score on day 3 and experienced no change through the course of the study. Interestingly, only the group treated with 54043-5 LALA-PG saw an improvement in disease state, with a decrease in clinical score on days 4 and 5, reaching the lowest score compared to other groups.

Lung samples from a set of mice from each group were collected at day 2 and 5 p.i. to assess their viral titers by quantification of viral RNA using qRT-PCR and performing infectivity assays in cells to determine the 50% tissue culture effective dose (TCID50) of virus in the tissues. The qRT-PCR results showed no statistically significant difference between mice therapeutically administered any of the antibodies compared to vehicle at days 2 and 5 p.i., all with an average lung viral titer in the range of 10^8^–10^11^ Geq/mL (**Figure S6D**). There was also no statistical significance observed between groups by TCID50 assay at day 2 p.i., with all values in the range of ∼10^7^–10^8^ TCID50 per gram of lung tissue (TCID50/g) (**Figure 6F**). However, on day 5 p.i., the group administered COV2-2381 had a TCID50 value at the limit of detection (∼10^2^ TCID50/g) compared to other antibodies and the vehicle, which had values ∼10^6^ TCID50/g. We also analyzed the lung tissue at this time point for the presence of pro-inflammatory (IL-1b, IL-6, IL-17, G-CSF and IFN-ɣ), anti-inflammatory (IL-12 (p40), IL-12 (p70), IL-10 and IL-13), Th1/Th2-associated (IL-3, IL-4 and GM-CSF) cytokines and chemokines (eotaxin, KC, MCP-1, RANTES and MIP-1b) **(Figure S6E-H)**. We did not detect consistent cytokine recruitment patterns between groups (**Figure S6E-G**). However, within the chemokine panel, levels of MCP-1 in COV2-2381-treated mice were significantly lower (∼1000 pg/mL) than the vehicle, the 54043-5, and the 54043-5 LALA-PG treated groups (∼10,000 pg/mL). Additionally, levels of RANTES for the 54043-5 LALA-PG treated group were significantly higher (∼14,000 pg/mL) than the COV2-2381 and the 54043-5 treated groups (∼2,500 pg/mL) (**Figure S6H**). The presence of these chemokines indicates that macrophages are being recruited to the site of infection and that recruitment may be increased in 54043-5 LALA-PG treated mice, although it is necessary to confirm these results through additional experiments (Schall, Bacon et al. 1990, Fuentes, Durham et al. 1995, Gunn, Nelken et al. 1997, Haberstroh, Stilo et al. 1998).

Histopathology analysis of lung tissues collected at day 2 p.i. revealed few differences among all the tested animal groups (54043-5, 54043-5 LALA-PG, DENV-2D22 and COV2-2381) (**Figure S6I-L**). There was a minimal decrease in incidence of mixed or mononuclear cell alveolar, bronchoalveolar, and/or interstitial inflammation in the DENV-2D22 group compared to all the other groups. Similarly, at day 5 p.i. there were minimal differences among the antibody-treated groups. Alveolar hyperplasia was not observed in the 54043-LALA-PG and DENV-2D22 treated groups compared to vehicle controls, while incidence of mononuclear cell vascular/perivascular inflammation was increased in the 54043-5 and the DENV-2D22 groups compared to vehicle controls. Bronchointerstitial pneumonia accompanied by vasculitis, bronchiolar hyperplasia and necrosis was observed in deceased mice at day 5 p.i., which is a common histopathological observation associated with SARS-CoV-2-induced inflammation in the lung tissues of infected mice.

## DISCUSSION

The continued emergence of pathogenic coronaviruses in humans underscores the need for interventions with broad reactivity. Here, we identified a panel of cross-reactive antibodies, highlighted by a pan-betacoronavirus antibody, 54043-5, that targets the S2 subunit and binds the spikes of multiple SARS-CoV-2 variants of concern as well as additional coronaviruses that infect other species. We show that 54043-5 targets a cryptic epitope at the apex of S2 that is inaccessible in the intact prefusion spike. Its elicitation by natural infection – either by SARS-CoV-2 or by previous infection with a seasonal coronavirus – may be a result of transient accessibility to the epitope during dynamic movement of the spike protein *in vivo*. Other class I fusion proteins such as influenza HA, VSV G, RSV F, and hMPV F have been observed to undergo trimeric “breathing”, granting access to epitopes at the trimeric interface (Albertini, Mérigoux et al. 2012, Bajic, Maron et al. 2019, Bangaru, Lang et al. 2019, Gilman, Furmanova-Hollenstein et al. 2019, Watanabe, McCarthy et al. 2019, Huang, Diaz and Mousa 2020, Simmons, Finney et al. 2023). Recently, the SARS-CoV-2 spike was also shown to undergo a reversible transition between an open and closed trimeric state, and binding of antibody 3A3, which binds a similar epitope to 54043-5 at the HR1/CH hinge, traps the spike in an asymmetric open trimer (Costello, Shoemaker et al. 2022). Further, 3D negative-stain EM reconstructions of a public class of S2 apex-directed antibodies isolated from convalescent patients revealed binding to a splayed SARS-CoV-2 S trimer with a preference for less stabilized prefusion spikes (Claireaux, Caniels et al. 2022). Our inability to observe complexes of 54043-5 bound to the soluble spike ectodomain by negative stain or cryo-EM indicates that antibody binding may similarly disrupt the closed, prefusion spike.

Although S2-directed antibodies are naturally elicited and common in SARS-CoV-2 convalescent repertoires most are weakly or non-neutralizing (Raybould, Kovaltsuk et al. 2021). Most of the S2-directed antibodies characterized to date target the membrane-proximal stem region and are broadly reactive, though most only moderately neutralize authentic and pseudotyped SARS-CoV-2 (Pinto, Sauer et al. 2021, Sauer, Tortorici et al. 2021, Wang, van Haperen et al. 2021, Li, Chao et al. 2022, Zhou, Yuan et al. 2022, Dacon, Peng et al. 2023, Zhou, Song et al. 2023) or fail to neutralize (Hsieh, Werner et al. 2021, Wang, van Haperen et al. 2021, Zhou, Yuan et al. 2022). More recently, antibodies have been isolated that bind the fusion peptide, which is located on the periphery of the S2 subunit (Dacon, Tucker et al. 2022, Low, Jerak et al. 2022, Sun, Yi et al. 2022, Dacon, Peng et al. 2023). The cryptic nature of these epitopes, and their impaired binding to stabilized 2P and 6P spikes indicate that binding of these antibodies also requires transient opening of the spike trimer. These antibodies neutralize pseudotyped and authentic SARS-CoV-2 more effectively than stem-helix-binding antibodies but are less potent than neutralizing RBD-directed antibodies. Antibodies that bind the S2 apex are less well characterized. Here, we observed no 54043-5-mediated neutralization of pseudotyped or authentic SARS-CoV-2 virus. Other S2 apex-binding antibodies, such as 3A3 and RAY53, incompletely neutralize SARS-CoV-2 pseudoviruses and fail to neutralize authentic virus, and S2 apex-directed IGHV1-69/IGKV3-11 antibodies were non-neutralizing in pseudovirus assays (Claireaux, Caniels et al. 2022, Silva, Huang et al. 2023). Together, these findings have implications for vaccine development strategies. Vaccines presenting S2-only antigens have been explored as a means to direct the immune response toward broadly reactive epitopes to minimize the likelihood of escape by future variants (Halfmann, Frey et al. 2022, Ng, Faulkner et al. 2022, Pang, Lu et al. 2022, Lee, Stewart et al. 2023). However, further investigation of the tradeoff between the reactive breadth and neutralization capacity of S2-directed antibodies and their effects on vaccine-induced immunity is necessary.

Non-neutralizing antibodies that target SARS-CoV-2 S dominate the repertoires of vaccinated individuals and comprise a substantial portion of the antibodies elicited by natural infection (Amanat, Thapa et al. 2021, Dugan, Stamper et al. 2021). Many of these antibodies induce Fc-mediated immune responses such as ADCP, ADCT, or ADCC that may confer protection *in vivo* (Shiakolas, Kramer et al. 2021). Although resistance to neutralizing sera has been documented for SARS-CoV-2 variants, including the recent Omicron variants, vaccine-induced antibodies maintain these Fc-mediated effector functions (Bartsch, Tong et al. 2022).

Further, breakthrough infections with Delta and Omicron variants led to an expansion of S2-directed, Fc receptor-binding antibodies (McNamara, Maron et al. 2022). Together, this indicates a substantial role for broadly reactive S2-directed antibodies – neutralizing and non-neutralizing – in strategic development of effective countermeasures for the ongoing threat from coronaviruses.

Here, we found that although wild-type 54043-5 can activate ADCP and ADCT *in vitro*, these functions do not translate into prophylactic or therapeutic protection in mice. Dose-dependence has been observed for antibody-mediated phagocytosis of spike-coated beads by THP-1 cells (Bahnan, Wrighton et al. 2021). When treated with varying concentrations of spike-binding antibodies, the percentage of THP-1 cell-spike interactions increased with dose but reached a threshold past which association began to decrease. This was also observed in mice treated therapeutically with a neutralizing, opsonic antibody after challenge with SARS-CoV-2 (Bahnan, Wrighton et al. 2021). Mice treated with 5x the protective dose of neutralizing antibody fared about as poorly as untreated mice. Though more experiments are needed to further explore the effect of dose on phagocytosis and protection, this may partly explain the discrepancy between our *in vitro* ADCP and protection data for the wild-type 54043-5 antibody. Phagocytosis is also affected by other properties that differ between antigens displayed on coated beads and those on the viral surface. As has been observed for HIV-1, low spike densities present on SARS-CoV-2 virions may prevent bivalent binding of IgGs, which is important for the promotion of phagocytosis (Klein and Bjorkman 2010, Ke, Oton et al. 2020). The height and angular position of spikes also vary on SARS-CoV-2 virions and can have implications for accessibility of the Fc to approaching phagocytes (Bakalar, Joffe et al. 2018, Ke, Oton et al. 2020). There are many additional factors that affect activation of ADCP *in vivo* that depend on the type of Fc receptor or phagocyte being engaged, the location and cell or tissue type, and related signaling pathways (Tay, Wiehe and Pollara 2019).

Although the set of LALA-PG substitutions introduced into the 54043-5 Fc region is one of the most effective at silencing Fc-mediated activity, very low levels of FcR binding have been reported for these antibodies (Wilkinson, Anderson et al. 2021). We show that in the lungs of infected mice treated with 54043-5 LALA-PG, the release of MCP-1 is maintained and the release of RANTES is increased, indicating that macrophages are being recruited to the lungs of infected mice. Further, although C1q binding was not detected by LALA-PG in murine IgG2A (analogous to IgG1 in humans), C3 fixation was reduced by just over 50%, indicating some complement activation can still occur (Lo, Kim et al. 2017). Overall, tight regulation of Fc-mediated immune responses is important for viral disease, but there remains much to learn. Our data indicate that the LALA-PG substitutions balance the effect of 54043-5 within these processes to confer protection in mice, though how this occurs is unknown. Treating mice with 54043-5 Fab could help begin to elucidate the effects of the Fc in disease progression. Additional experiments that test T-cell activation and complement fixation by wild-type and LALA-PG 54043-5 antibodies would also help to better understand the effects of Fc attenuation.

## Acknowledgements

We thank A. Jones, L. Raju and J. Roberson of VANTAGE for their expertise regarding next-generation sequencing and library preparation, D. Flaherty and B. Matlock of the Vanderbilt Flow Cytometry Shared Resource for help with flow panel optimization, and members of the Georgiev laboratory for comments on the manuscript. The Vanderbilt VANTAGE Core provided technical assistance for this work. VANTAGE is supported, in part, by CTSA grant 5UL1 RR024975-03, the Vanderbilt Ingram Cancer Center (P30 CA68485), the Vanderbilt Vision Center (P30 EY08126) and NIH/NCRR (G20 RR030956). Flow cytometry experiments were performed in the Vanderbilt University Medical Center (VUMC) Flow Cytometry Shared Resource. The VUMC Flow Cytometry Shared Resource is supported by the Vanderbilt Ingram Cancer Center (P30 CA68485) and the Vanderbilt Digestive Disease Research Center (DK058404). Vanderbilt University Medical Center has utilized the non-clinical and pre-clinical services program offered by the National Institute of Allergy and Infectious Diseases. The following reagent was obtained through BEI Resources, NIAID, NIH: Human Embryonic Kidney Cells (HEK-293T) Expressing Human Angiotensin-Converting Enzyme 2, HEK-293T-hACE2 Cell Line, NR-52511. For work described in this manuscript, I.S.G., K.J.K, S.C.W., A.R.S., K.A.P., A.A.A., and C.M.H. were supported, in part, by NIH National Institute of Allergy and Infectious Diseases (NIAID) award R01 AI131722-S1, the Hays Foundation COVID-19 Research Fund, Fast Grants, and the G. Harold and Leila Y. Mathers Charitable Foundation. J.S.M. and N.V.J. were supported, in part, by NIAID award R01 AI131722-S1 and by Welch Foundation grant no. F-0003-19620604. J.E.C., N.S., R.E.S., R.S.N., and R.H.C. were supported, in part, by Defense Advanced Research Projects Agency (DARPA) grant HR0011-18-2-0001, US NIH contract 75N93019C00074, NIH grant R01 AI157155, the Dolly Parton COVID-19 Research Fund at Vanderbilt and a grant from Fast Grants, Mercatus Center, George Mason University. J.E.C. is a recipient of the 2019 Future Insight Prize from Merck KGaA, which supported this work with a grant.

## Author Contributions

N.V.J., S.C.W., K.J.K., I.S.G., and J.S.M. developed the methodology.

N.V.J., S.C.W., K.J.K., C.M.H., S.P., S.I.R., N.S., E.A., I.P., G.P., G.P., Y.H., P.G., J.D.A., N.U.,

A.R.S., K.A.P., R.S.N., R.E.S., A.A.A., and R.P. performed the investigations. N.V.J., S.C.W., and K.J.K. performed validations. N.V.J., S.C.W., K.J.K., C.M.H., J.S.M., and I.S.G. wrote the original draft. All authors reviewed and edited the manuscript. J.S.M. and I.S.G. acquired funding. J.S.M., I.S.G., J.E.C., A.B., T.M.R., B.F.H., G.A.S., and R.H.C. provided resources. J.S.M. and I.S.G. supervised the work.

## EXPERIMENTAL MODELS AND SUBJECT DETAILS

### Human Subjects

(For samples 54041, 54042, and 54043) The 45-year-old, male donor had previous laboratory-confirmed COVID-19, 3 months prior to blood collection. The donor for sample 54044 was a healthy 45-year-old adult male. No other information on this donor is known. The studies were reviewed and approved by the Institutional Review Board of Vanderbilt University Medical Center. The samples were obtained after written informed consent was obtained.

### Cell Lines

A variety of cell lines were used for different assays in this study. Expi293F mammalian cells (ThermoFisher, A14527) were maintained in FreeStyle F17 expression medium supplemented with a final concentration of 0.1% Pluronic Acid F-68 and 4mM L-Glutamine. ExpiCHO cells (ThermoFisher, A29127) were maintained in ExpiCHO Expression medium (ThermoFisher, A2910002). Cells were cultured at 37 °C with 8% CO2 saturation while shaking. Vero E6 cells (ATCC, CRL-1586) and all HEK293T cell lines (ATCC, CRL-3216; ATCC, CRL-11268; BEI, NR-52511) were maintained in Dulbecco’s minimal essential medium (DMEM) supplemented with 10mM HEPES pH 7.3, 1X non-essential amino acids, 1mM sodium pyruvate, 100U/mL of penicillin-streptomycin, and 10% fetal bovine serum and grown in 37 C with 5% CO2. Authentication analysis was not performed on the cell lines used. Jurkat-Lucia NFAT-CD16 cells were maintained in IMDM media with 10% heat-inactivated fetal bovine serum (Gibco, Gaithersburg, MD), 1% Penicillin Streptomycin (Gibco, Gaithersburg, MD) and 10 mg/mL of Blasticidin and 100 mg/mL of Zeocin was added to the growth medium every other passage. THP-1 cells were used for both the ADCP and ADCT assays and obtained from the AIDS Reagent Program, Division of AIDS, NIAID, NIH contributed by Dr. Li Wu and Vineet N. Kewal Ramani. Cells were cultured at 37 °C, 5% CO2 in RPMI containing 10% heat-inactivated fetal bovine serum (Gibco, Gaithersburg, MD) with 1% Penicillin Streptomycin (Gibco, Gaithersburg, MD) and 2-mercaptoethanol to a final concentration of 0.05 mM and not allowed to exceed 4 3 105 cells/mL to prevent differentiation.

### Viruses

The generation of a replication-competent VSV expressing SARS-CoV-2 S protein with a 21 amino-acid C-terminal deletion that replaces the VSV G protein (VSV-SARS-CoV-2) was described previously (**Case et al., 2020b**). The S protein-expressing VSV virus was propagated in MA104 cell culture monolayers (African green monkey, ATCC CRL-2378.1) as described previously (**Case et al., 2020b**), and viral stocks were titrated on Vero E6 cell monolayer cultures. VSV plaques were visualized using neutral red staining. All work with infectious SARS-CoV-2 was performed in Institutional Biosafety Committee approved BSL3 and A-BSL3 facilities at Washington University School of Medicine using appropriate positive pressure air respirators and protective equipment.

## METHOD DETAILS

### Antigen expression and purification

An assortment of recombinant soluble protein antigens was used in the LIBRA-seq experiment and assays. All Expi293F cells were cultured at 8% CO2 saturation and 37°C with shaking in FreeStyle F17 expression media (Thermo Fisher) supplemented to a final concentration of 0.1% Pluronic Acid F-68 and 4 mM L-glutamine.

Plasmids were transiently transfected in Expi293F cells using polyethylenimine or ExpiFectamine™ transfection reagent (Thermo Fisher Scientific) and encoded the following: residues 1–1208 of the SARS-CoV-2 spike with a mutated S1/S2 cleavage site, proline substitutions at positions 817, 892, 899, 942, 986 and 987, and a C-terminal T4-fibritin trimerization motif, an 8x HisTag, and a TwinStrepTag (SARS-CoV-2 S Hexapro (HP)); 1–1208 of the SARS-CoV-2 spike with a mutated S1/S2 cleavage site, proline substitutions at positions 817, 892, 899, 942, 986 and 987, as well as mutations L18F, D80A, L242-244L del, R246I, K417N, E484K, N501Y, and a C-terminal T4-fibritin trimerization motif, an 8x HisTag, and a TwinStrepTag (SARS-CoV-2 spike HP Beta); 1–1208 of the SARS-CoV-2 spike with a mutated S1/S2 cleavage site, proline substitutions at positions 817, 892, 899, 942, 986 and 987, as well as mutations 69-70del, Y144del, N501Y, A570D, P681H, and a C-terminal T4-fibritin trimerization motif, an 8x HisTag, and a TwinStrepTag (SARS-CoV-2 spike HP Alpha); residues 1-1190 of the SARS-CoV spike with proline substitutions at positions 968 and 969, and a C-terminal T4-fibritin trimerization motif, an 8x HisTag, and a TwinStrepTag (SARS-CoV S-2P); residues 1-1291 of the MERS-CoV spike with a mutated S1/S2 cleavage site, proline substitutions at positions 1060 and 1061, and a C-terminal T4-fibritin trimerization motif, an AviTag, an 8x HisTag, and a TwinStrepTag (MERS-CoV S-2P Avi); residues 1-1278 of the HCoV-OC43 spike with proline substitutions at positions 1070 and 1071, and a C-terminal T4-fibritin trimerization motif, an 8x HisTag, and a TwinStrepTag (HCoV-OC43 S-2P); residues 1-1277 of the HCoV-HKU1 spike with a mutated S1/S2 cleavage site, proline substitutions at positions 1067 and 1068, and a C-terminal T4-fibritin trimerization motif, an 8x HisTag, and a TwinStrepTag (HCoV-HKU1 S-2P); 1–1208 of the SARS-CoV-2 spike with a mutated S1/S2 cleavage site, proline substitutions at positions 817, 892, 899, 942, 986 and 987, as well as mutations T19R, del157/158, L452R, T478K, D614G, P681R, D950N, and a C-terminal T4-fibritin trimerization motif, Avitag, HRV3C, 8x HisTag, and a TwinStrepTag (SARS-CoV-2 Delta S HP); 1–1208 of the SARS-CoV-2 spike with a mutated S1/S2 cleavage site, proline substitutions at positions 817, 892, 899, 942, 986 and 987, as well as mutations A67V, del69/70, T95I, G142D, del143/145, del11, L212I, G339D, S371L, S373P, S375F, S477N, T478K, E484A, Q493R, Q496S, Q498R, N501Y, Y505H, T547K, D614G, H655Y, N679K, P681H, N764K, D796Y, N856K, Q954H, N969K, L981F, and a C-terminal T4-fibritin trimerization motif, Avitag, HRV3C, 8x HisTag, and a TwinStrepTag (SARS-CoV-2 Omicron BA.1 S HP); 1–1208 of the SARS-CoV-2 spike with a mutated S1/S2 cleavage site, proline substitutions at positions 817, 892, 899, 942, 986 and 987, as well as mutations T19I, Del24-26, G142D, V213G, G339D, S371F, S373P, S375F, T376A, D405N, R408S, K417N, N440K, S477N, T478K, E484A, Q493R, Q498R, N501Y, Y505H, D614G, H655Y, N679K, P681H, N764K, D796Y, Q954H, N969K, and a C-terminal T4-fibritin trimerization motif, Avitag, HRV3C, 8x HisTag, and a TwinStrepTag (SARS-CoV-2 Omicron BA.2 S HP); 1–1383 of the TGEV spike (Genbank: P07946.2) with proline substitutions at positions 1139-1140 (E1139P, L1140P), and a C-terminal T4-fibritin trimerization motif, an 8x HisTag, and a TwinStrepTag (TGEV-2P)). 1–1093 of the SDCV spike (Genbank: AMN91621.1) with proline substitutions at positions 855-856 (E855P, V856P), and a C-terminal T4-fibritin trimerization motif, an 8x HisTag, and a TwinStrepTag (SDCV-2P)). 1–1191 of the WIV1 spike (Genbank: AGZ48828.1) with proline substitutions at positions 969-970 (K969P, V970P), and a C-terminal T4-fibritin trimerization motif, an 8x HisTag, and a TwinStrepTag (WIV1-2P)). 1–1319 of the PEDV spike (Genbank: NP_598310.1) with proline substitutions at positions 1073-1074 (I1073P, L1074P), and a C-terminal T4-fibritin trimerization motif, an 8x HisTag, and a TwinStrepTag (PEDV-2P)). 1–1207 of the HKU9 spike (Genbank: YP_001039971.1) with proline substitutions at positions 983-984 (G983P, L984P), and a C-terminal T4-fibritin trimerization motif, an Avitag,, an HRV 3C protease site, an 8x HisTag, and a TwinStrepTag (HKU9-2P)). All coronavirus spike supernatants were collected 5-7 days post transfection, sterile filtered, and purified over a StrepTrap XT column (Cytiva Life Sciences). Purified proteins were further purified using a size exclusion Superose6 Increase column (Cytiva Life Sciences). For LIBRA-seq, the purified antigens were then biotinylated with the EZ-Link Sulfo-NHS-Biotin (Thermo Fisher Scientific) using a 50:1 biotin to protein molar ratio for calculations.

Recombinant ACE2 ectodomain (Genbank: BAJ21180.1) with the addition of an 8x HisTag and a StrepTag II was expressed and purified in the same manner as the CoV spike antigens.

Recombinant HIV-1 gp140 SOSIP trimer from strain ZM197 (clade C)(Georgiev, Joyce et al. 2015) containing an AviTag was cultured in Expi293F cells and transfected in the same method as above. The clarified supernatant was run over an affinity column of agarose-bound *Galanthus nivalis* lectin (GNA, Snowdrop) slowly at 4°C. The column was washed with 1X PBS and bound protein was eluted with 1M methyl-a-D-mannopyranoside in PBS. The protein eluate was buffer exchanged into 1X PBS and then purified by size exclusion chromatography using a Superdex 200 Increase 10/300 GL Sizing column on the AKTA FPLC system (GE Life Sciences). The fractions of purified protein were analyzed by SDS-PAGE and binding was confirmed using ELISA with known antibodies.

The recombinant HA proteins (A/New Caledonia/20/99 H1N1 GenBank ACF41878 (NC99) and A/California/07/2009 H1N1 Genbank FJ969540.1) was produced using Expi 293F cells and the Expifectamine 293 transfection reagent. The protein contains the HA ectodomain with a point mutation at the sialic acid-binding site (Y98F), T4 fibritin foldon trimerization domain, AviTag, and hexahistidine-tag. The cells were cultured for 4-5 days, then the supernatant was harvested and sterile filtered. The pH and NaCl concentration were adjusted by adding 1M Tris-HCl (pH 7.5) and 5M NaCl to 50 mM and 500 mM, respectively. The supernatant was then mixed with Ni Sepharose excel resin (GE Healthcare) to capture the hexahistidine tag. The resin was isolated in a column by gravity and the captured HA protein was eluted by a Tris-NaCl (pH 7.5) buffer containing 300 mM imidazole. The eluted protein was further purified by size exclusion chromatography using a HiLoad 16/60 Superdex 200 column (GE Healthcare). Fractions containing the appropriate sized HA protein were concentrated, analyzed by SDS-PAGE, and tested for antigenicity by ELISA using known antibodies. The proteins were then stored at -80C until use.

The SARS-CoV-2 S1, SARS-CoV-2 S2, SARS-CoV-2 RBD, SARS-CoV-2 NTD, MERS-CoV S1, and MERS-CoV S2 subdomains as well as recombinant HCoV-NL63 and HCoV-229E S were purchased from Sino Biological.

### Oligonucleotide barcodes

We used oligos that possess 15 bp antigen barcode, a sequence capable of annealing to the template switch oligo that is part of the 10X bead-delivered oligos and contain truncated TruSeq small RNA read 1 sequences in the following structure: 50CCTTGGCACCCGAGAATTCCANNNNNNNNNNNNNCCCATATAAGA*A*A-30, where Ns represent the antigen barcode. We used the following antigen barcodes: GCAGCGTATAAGTCA (SARS-CoV-2 S), AACCCACCGTTGTTA (SARS-CoV-2 S D614G), GCTCCTTTACACGTA (SARS-CoV S), GGTAGCCCTAGAGTA (MERS-CoV S), AGACTAATAGCTGAC (HCoV-OC43 S), TGTGTATTCCCTTGT (HCoV-HKU1) GACAAGTGATCTGCA (HCoV-NL63 S), GTGTGTTGTCCTATG (HCoV-229E S), TACGCCTATAACTTG (ZM197 HIV EnV), TCATTTCCTCCGATT (HA NC99), TGGTAACGACAGTCC (SARS-CoV RBD-SD1), TTTCAACGCCCTTTC (SARSCoV-2 RBD-SD1), GTAAGACGCCTATGC (MERS-CoV RBD), CAGTAAGTTCGGGAC (SARS-CoV-2 NTD), Oligos were ordered from IDT with a 5’ amino modification and HPLC purified.

### Labeling antigens with DNA oligonucleotide barcodes

For each antigen described above, the unique DNA barcodes were directly conjugated to the antigen using a SoluLINK Protein-Oligonucleotide Conjugation kit (TriLink, S-9011) according to kit protocol. In short, we desalted the oligonucleotide and protein, modified the amino-oligonucleotide with the 4FB cross-linker, and modified the biotinylated antigen with S-HyNic. Afterwards, the 4FB-oligonucleotide and the HyNic-antigen were mixed to form a stable bond between the protein and the oligonucleotide. The antigen-oligonucleotide concentrations were then determined using a bicinchoninic acid (BCA) assay, and the HyNic molar substitution ratios of each antigen-oligonucleotide conjugate was determined using a NanoDrop according to SoluLINK protocol instructions. Excess oligonucleotide was removed from the protein-oligonucleotide conjugates using an AKTA FPLC and were subsequently verified using SDS-PAGE and silver stain. The optimal amounts of antigen-oligonucleotide conjugates to be used in antigen-specific B cell sorting were then determined through flow cytometry titration experiments on cell lines expressing BCRs of known specificities.

### Antigen specific B cell sorting

To start, PBMCs were thawed, washed, and counted. Viability was evaluated using Trypan Blue. The cells were then washed with a solution of DPBS supplemented with 0.1% Bovine serum albumin (BSA). Afterwards, the cells were resuspended in DPBS-BSA and stained with cell markers: Ghost Red 780 for viability, CD14-APC-Cy7, CD3-FITC, CD19-BV711 and IgG-PE-Cy5. Additionally, antigen-oligo conjugates were added to the stain. After a 30-minute incubation in the dark at room temperature, the cells were washed again three times with DPBS-BSA at 300 g for five minutes. Then, the cells were incubated for 15 minutes at room temperature with Streptavidin-PE to label cells with bound antigen. The cells were again washed three times with DPBS-BSA, resuspended in DPBS, and sorted by FACS. Antigen positive B cells were bulk sorted and delivered to the Vanderbilt Technologies for Advanced Genomics (VANTAGE) sequencing core at an appropriate target concentration for 10X Genomics library preparation and subsequent sequencing. FACS data were analyzed using FlowJo.

### Sample and library preparation for sequencing

Single-cell suspensions were processed using the Chromium Controller microfluidics device (10X Genomics) and the B cell Single Cell V(D)J solution as per the manufacturer’s instructions. The aim was to capture 10,000 B cells per 1/8 10X cassette. Slight modifications were made to intercept, amplify, and purify the antigen barcode libraries, as previously described (Setliff, Shiakolas et al. 2019).

### Sequence processing and bioinformatics analysis

Our established pipeline was followed, which takes paired-end FASTQ files of oligonucleotide libraries as input, to process and annotate reads for cell barcodes, unique molecular identifiers (UMIs) and antigen barcodes, resulting in a cell barcode-antigen barcode UMI count matrix (Setliff, Shiakolas et al. 2019). B cell receptor contigs were processed using CellRanger (10x Genomics) and GRCh38 as reference, while the antigen barcode libraries were also processed using CellRanger (10x Genomics). The cell barcodes that overlapped between the two libraries formed the basis of the subsequent analysis. Cell barcodes that had only non-functional heavy chain sequences as well as cells with multiple functional heavy chain sequences and/or multiple functional light chain sequences, were eliminated, reasoning that these may be multiplets. We also aligned the B cell receptor contigs (filtered_contigs.fasta file output by CellRanger, 10x Genomics) to IMGT reference genes using HighV-Quest (Alamyar, Duroux et al. 2012). The output of HighV-Quest was parsed using ChangeO (Gupta, Vander Heiden et al. 2015), and combined with an antigen barcode UMI count matrix. Finally, we determined the LIBRA-seq score for each antigen in the library for every cell as previously described (Setliff, Shiakolas et al. 2019).

### High-throughput antibody microscale expression and purification

For high-throughput production of recombinant antibodies, a microscale approach was employed. For antibody expression, microscale transfection was performed (∼1 ml per antibody) with CHO cell cultures using the Gibco ExpiCHO Expression System and a protocol for deep 96-well blocks (Thermo Fisher Scientific). Briefly, synthesized antibody-encoding DNA (∼2 μg per transfection) was added to OptiPro serum-free medium (OptiPro SFM), incubated with ExpiFectamine CHO Reagent, and added to 800 µl of ExpiCHO cell cultures in deep 96-well blocks using a ViaFlo384 liquid handler (Integra Biosciences). The plates were incubated on an orbital shaker at 1,000 r.p.m. with an orbital diameter of 3 mm at 37 °C in 8% CO2. The next day after transfection, ExpiFectamine CHO Enhancer and ExpiCHO Feed reagents (Thermo Fisher Scientific) were added to the cells, followed by 4 more days of incubation. Culture supernatants were collected after centrifuging the blocks at 450 g for 5 minutes and were stored at 4°C until use. For high-throughput microscale antibody purification, fritted deep-well plates were used containing 25 μl of settled Protein G resin (GE Healthcare Life Sciences) per well. Clarified culture supernatants were incubated with protein G resin for antibody capturing, washed with PBS using a 96-well plate manifold base (Qiagen) connected to the vacuum and eluted into 96-well PCR plates using 86 μl of 0.1 M glycine-HCl buffer, pH 2.7. Purified antibodies were then neutralized with 14 μl of 1 M Tris-HCl pH 8.0 and buffer exchanged into PBS using Zeba Spin Desalting plates (Thermo Fisher Scientific). Purified antibodies were then stored at 4°C until use.

### Antibody expression and purification

Variable heavy and light genes were inserted into custom plasmids that encode the constant region for the human IgG1 heavy chain and respective lambda and kappa light chains (pTwist CMV BetaGlobin WPRE Neo vector, Twist Bioscience). The antibodies were expressed in Expi293F cells by co-transfecting heavy chain and light chain expressing plasmids using polyethylenimine or Expifectamine transfection reagent, and the cells were cultured for 4-5 days. These cells were maintained as previously described in the antigen purification methods. Cultures were harvested, centrifuged and supernatant was 0.45 µm filtered with Nalgene Rapid Flow Disposable Filter Units with PES membrane. The filtered supernatant was run over a column containing Protein A agarose resin that was equilibrated with PBS. The column was washed with PBS, and then the antibodies were eluted with 100mM Glycine HCl at 2.7 pH directly into a 1:10 volume of 1M Tris-HCl pH 8.0. Eluted antibodies were buffer exchanged into PBS using using Amicon Ultra centrifugal filter units, centrifuging and topping off three times with PBS, and finally concentrated. The antibodies were analyzed by SDS-PAGE. Antibody plasmids were sequenced to confirm the expected heavy and light chain match.

## ELISA

To evaluate the binding of the expressed antibodies, soluble purified antigen was plated at a concentration of 2 µg/mL and incubated overnight at 4°C. The next day, the plates were washed three times with a PBS solution containing 0.05% Tween-20 (PBS-T) and then coated with 5% milk powder in PBS-T. The plates were incubated for one hour at room temperature and then washed three times with PBS-T. The primary antibodies were diluted in 1% milk in PBS-T, starting at a concentration of 10 µg/mL with a serial 1:5 or 1:10 dilution and then added to the plate. The plates were incubated for an additional hour at room temperature and then washed three times with PBS-T. The secondary antibody, goat anti-human IgG conjugated to peroxidase, was added at a dilution of 1:10,000 in 1% milk in PBS-T to the plates, which were incubated for one hour at room temperature. The plates were washed three times with PBS-T and then developed by adding TMB substrate to the plates. The plates were incubated for ten minutes at room temperature, and the reaction was stopped with 1N sulfuric acid. Plates were read at 450 nm. The data is shown as one representative biological replicate with the mean ± SEM for one ELISA experiment. The ELISAs were repeated 2 or more times. The area under the curve (AUC) was calculated using GraphPad Prism 9.5.0.

### Autoreactivity

Monoclonal antibody reactivity to nine autoantigens (SSA/Ro, SS-B/La, Sm, ribonucleoprotein (RNP), Scl 70, Jo-1, dsDNA, centromere B, and histone) was measured using the AtheNA Multi-Lyte® ANA-II Plus test kit (Zeus scientific, Inc, #A21101). Antibodies were incubated with AtheNA beads for 30min at concentrations of 50, 25, 12.5 and 6.25 µg/mL. Beads were washed, and incubated with secondary and read on the Luminex platform as specified in the kit protocol. Data were analyzed using AtheNA software. Positive (+) specimens received a score > 120, and negative (-) specimens received a score < 100. Samples between 100-120 were considered indeterminate.

### Surface Plasmon Resonance

His-tagged SARS CoV-2 S2 (S2-37) was immobilized to a single flow cell of an NTA sensorchip a Biacore X100 (GE Healthcare). Two samples containing only running buffer (10 mM HEPES pH 8.0, 150 mM NaCl and 0.005% Tween 20) were injected over the ligand and reference flow cells, followed by injection of Fab 54043-5 serially diluted from 64-0.25 nM. The chip was regenerated between cycles using 0.35 M EDTA and 0.1 M NaOH followed by 0.5 mM NiCl2. The resulting data were double-reference subtracted and fit to a 1:1 binding model using the Biacore X100 Evaluation software.

### EM sample prep and data collection

SARS-CoV-2 S2 was expressed and purified as in (CLH ref). To form the S2-Fab 54043-5 complex for cryo-EM studies, purified SARS-CoV-2 S2 and Fab 54043-5 were combined at final concentrations of 0.75 mg/mL and 0.3 mg/mL respectively in buffer containing 2 mM Tris pH 7.5, 200 mM NaCl, 0.02% NaN3. 3µL of sample was applied to Au-300 1.2/1.3 grids (UltrAuFoil) that had been plasma cleaned for 4 minutes using a 4:1 ratio of O2:H2 in a Solarus 950 plasma cleaner (Gatan). Grid freezing was carried out using a Vitrobot Mark IV (ThermoFisher) set to 100% humidity and 4 C°. Excess liquid was blotted from the grids using a blot force of 0 for 4 seconds after a 10 second wait and immediately plunge-frozen in liquid ethane. 1,885 movies were collected from a single grid in a Talos F200C (Thermo Fisher) equipped with a Falcon 4 detector (Thermo Fisher). All movies were collected using SerialEM automation software (Mastronarde 2005). Particles were imaged at a calibrated magnification of 0.94 Å/pixel, with a dose of 6 e-/pix/sec for 8 seconds for a total dose of 50 e/Å2. Additional details about data collection parameters can be found in **Table S2**.

### Cryo-EM

Motion correction, CTF estimation, particle picking, and preliminary 2D classification were performed using cryoSPARC v3.3.1 live processing (Punjani, Rubinstein et al. 2017). Particles were initially extracted with a box size of 532 pixels with Fourier cropping to a box size of 160 pixels. The final iteration of 2D class averaging distributed 793,124 particles into 50 classes using an uncertainty factor of 1. From that, 218,540 particles were selected and used to perform an *Ab inito* reconstruction with four classes followed by heterogeneous refinement of those four classes using all selected particles. 127,773 particles combined from the two highest quality classes – one with 3 copies of Fab 54043-5 bound to S2 and one that appeared to be S2 alone – were used for homogenous refinement of the Fab-bound volume with no applied symmetry. After a single round of homogenous refinement, particles were re-extracted using an uncropped box size of 532 pixels and duplicate particles were removed. The remaining 126,621 particles were used to perform a non-uniform refinement of the previous volume without applied symmetry, followed by a non-uniform refinement with applied C3 symmetry and with optimized per-group CTF parameters enabled to yield a final 3.0 Å volume (Rubinstein and Brubaker 2015). To improve map quality, the final volume was processed using the DeepEMhancer tool via COSMIC^2^ science gateway (Sanchez-Garcia, Gomez-Blanco et al. 2021). An initial model was generated by docking the S2 subunit within PDBID: 6XKL (residues 703-1147) and a Fab model generated based on the 54043-5 sequence using SAbPred ABodyBuilder into the refined volume via ChimeraX (Dunbar, Krawczyk et al. 2016, Goddard, Huang et al. 2018, Stewart-Jones, Chuang et al. 2018, Pettersen, Goddard et al. 2021, Meng, Goddard et al. 2023). The model was iteratively refined and completed using a combination of Phenix, Coot, and ISOLDE (Adams, Grosse-Kunstleve et al. 2002, Emsley and Cowtan 2004, Croll 2018). The full cryo-EM processing workflow and structure validation can be found in **Supplementary Figures S2 and S3 and Table S2**.

### Conservation Analysis

*Orthocoronavirinae* species and strains were taken from the International Committee on Taxonomy of Viruses (Lefkowitz, Dempsey et al. 2018). In figure panels 2A and 4C-D, one strain from each viral species in the *Betacoronavirus* genus was used with the exception of using two highly relevant *Betacoronavirus 1*, and four *SARS-related CoV* strains. Figure 2A was generated in MEGA (Tamura, Stecher and Kumar 2021) using the built-in Maximum Likelihood method and 4C was generated using the MUSCLE algorithm in MEGA. For figure panel 4D, the sequence alignment generated for panel 4C was imported into ChimeraX (Goddard, Huang et al. 2018, Pettersen, Goddard et al. 2021, Meng, Goddard et al. 2023) to map the sequence conservation onto the model of the SARS-CoV-2 spike S2 subunit. Sequence-based structural conservation was calculated within ChimeraX using the entropy-based measure from AL2CO (Pei and Grishin 2001) using a range of -1.4 – 1.4, illustrated as a color gradient from white to purple. Figure 4E was calculated taking one representative strain from each *Orthocoronavirinae* species and calculating the percent of strains within a genus with the same residue at a given spike position as that of SARS-CoV-2.

### Buried Solvent Accessible Surface Area Analysis

Total buried solvent accessible surface area (BSA) of 54043-5, just the heavy chain, or just the light chain was calculated with PDBePISA (Krissinel and Henrick 2007). For residue-level BSA analysis, antibody-bound and unbound structures of spike were first prepared in PyMOL (Schrodinger 2015). The solvent accessible surface area (SASA) for each residue was then calculated with the DSSP (Kabsch and Sander 1983) tool and BSA was calculated as SASAunbound - SASAbound. This was performed for all coronavirus antibody-antigen structures listed on SAbDab (Dunbar, Krawczyk et al. 2014) as of the cutoff date February 20, 2023. These were filtered down to two primary S2-directed antibody groups: stem-helix and fusion-peptide antibodies. The numbering for all coronavirus S2 epitopes was converted to the SARS-CoV-2 spike numbering system, the BSA values were summed for each residue, and cells were shaded from white (0 BSA) to dark green (the highest BSA within a given row of Figure 4E).

### Public Clonotype Analysis

Public clonotype analysis was performed using all human SARS-CoV-2 spike-specific antibodies on the CoVAbDab (Raybould, Kovaltsuk et al. 2021) with the cutoff date of February 21, 2023. The calculations were performed using the V-genes and CDR3s taken directly from the database, with the exception of treating V-gene paralogues as the same gene.

### Plaque reduction neutralization test (PRNT)

The virus neutralization with live authentic SARS-CoV-2 virus was performed in the BSL-3 facility of the Galveston National Laboratory using Vero E6 cells (ATCC CRL-1586) following the standard procedure. Briefly, Vero E6 cells were cultured in 96-well plates (10^4^ cells/well). Next day, 4-fold serial dilutions of antibodies were made using MEM-2% FBS, as to get an initial concentration of 100 µg/ ml. Equal volume of diluted antibodies (60 µl) were mixed gently with authentic virus (60 µl containing 200 pfu) and incubated for 1 h at 37°C/5%CO2 atmosphere. The virus-serum mixture (100 µl) was added to cell monolayer in duplicates and incubated for 1 at h 37°C/ 5% CO2 atmosphere. Later, virus-serum mixture was discarded gently, and cell monolayer was overlaid with 0.6% methylcellulose and incubated for 2 days. The overlay was removed, and the plates were fixed in 4% paraformaldehyde twice following BSL-3 protocol. The plates were stained with 1% crystal violet and virus-induced plaques were counted. The percent neutralization and/or NT50 of antibody was calculated by dividing the plaques counted at each dilution with plaques of virus-only control. For antibodies, the inhibitory concentration at 50% (IC50) values were calculated in GraphPad Prism software by plotting the midway point between the upper and lower plateaus of the neutralization curve among dilutions. The Alpha variant virus incorporates the following substitutions: Del 69-70, Del 144, E484K, N501Y, A570D, D614G, P681H, T716I, S982A, D1118H. The Beta variant incorporates the following substitutions: Del 24, Del 242-243, D80A, D215G, K417N, E484K, N501Y, D614G, H665Y, T1027I. The Delta variant incorporates the following substitutions: T19R, G142D, Del 156-157, R158G, L452R, T478K, D614G, P681R, Del 689-691, D950N; the deletion at positions 689-691 has not been observed in nature, and was identified upon one passage of the virus.

### Real-time cell analysis (RTCA) neutralization assay

To determine neutralizing activity of IgG proteins, we used real-time cell analysis (RTCA) assay on an xCELLigence RTCA MP Analyzer (ACEA Biosciences Inc.) that measures virus-induced cytopathic effect (CPE) (Suryadevara, Gilchuk et al. 2022). Briefly, 50 μL of cell culture medium (DMEM supplemented with 2% FBS) was added to each well of a 96-well E-plate using a ViaFlo384 liquid handler (Integra Biosciences) to obtain background reading. A suspension of 18,000 Vero-E6 cells in 50 μL of cell culture medium was seeded in each well, and the plate was placed on the analyzer. Measurements were taken automatically every 15 min, and the sensograms were visualized using RTCA software version 2.1.0 (ACEA Biosciences Inc). VSV-SARS-CoV-2 (0.01 MOI, ∼120 PFU per well) was mixed 1:1 with a dilution of mAb in a total volume of 100 μL using DMEM supplemented with 2% FBS as a diluent and incubated for 1 h at 37°C in 5% CO2. At 16 h after seeding the cells, the virus-mAb mixtures were added in replicates to the cells in 96-well E-plates. Triplicate wells containing virus only (maximal CPE in the absence of mAb) and wells containing only Vero cells in medium (no-CPE wells) were included as controls. Plates were measured continuously (every 15 min) for 48 h to assess virus neutralization. Normalized cellular index (CI) values at the endpoint (48 h after incubation with the virus) were determined using the RTCA software version 2.1.0 (ACEA Biosciences Inc.). Results are expressed as percent neutralization in a presence of respective mAb relative to control wells with no CPE minus CI values from control wells with maximum CPE. RTCA IC50 values were determined by nonlinear regression analysis using Prism software.

### Antibody-dependent cellular phagocytosis (ADCP) assay

SARS-CoV-2 original or Beta spike was biotinylated using EZ link Sulfo-NHS-LC-Biotin kit (ThermoFisher) and coated on to fluorescent neutravidin beads as previously described (Ackerman, Moldt et al. 2011). Briefly, beads were incubated for two hours with monoclonal antibodies at a starting concentration of 2 µg/mL and titrated five-fold or plasma at a single 1 in 100 dilution. Opsonized beads were incubated with the monocytic THP-1 cell line overnight, fixed and interrogated on the FACSAria II. Phagocytosis score was calculated as the percentage of THP-1 cells that engulfed fluorescent beads multiplied by the geometric mean fluorescence intensity of the population less the no antibody control. For this and all subsequent Fc effector assays, pooled plasma from 5 PCR-confirmed SARS-CoV-2 infected individuals and CR3022 were used as positive controls and plasma from 5 pre-pandemic healthy controls and Palivizumab were used as negative controls. In addition, samples from both waves were run head-to-head in the same experiment. ADCP scores for original and Beta spikes were normalised to each other and between runs using CR3022.

### Antibody-dependent cellular cytotoxicity (ADCC) assay

The ability of plasma antibodies to cross-link and signal through FcgRIIIa (CD16) and spike expressing cells or SARS-CoV-2 protein was measured as a proxy for ADCC. For spike assays, HEK293T cells were transfected with 5µg of SARS-CoV-2 original variant spike (D614G), Beta, Gamma, Delta or SARS-1 spike plasmids using PEI-MAX 40,000 (Polysciences) and incubated for 2 days at 37 °C. Expression of spike was confirmed by differential binding of CR3022 and P2B-2F6 and their detection by anti-IgG APC staining measured by flow cytometry. For original or Beta NTD or RBD assays protein was coated at 1 mg/mL on a high binding ELISA 96-well plate and incubated at 4 °C overnight. Plates were then washed with PBS and blocked at room temperature for 1 hr with PBS + 2.5% BSA. Subsequently, protein or 1 x 10^5^ spike transfected cells per well were incubated with heat inactivated plasma (1:100 final dilution) or monoclonal antibodies (final concentration of 100 µg/mL) in RPMI 1640 media supplemented with 10% FBS 1% Pen/Strep (Gibco, Gaithersburg, MD) for 1 hour at 37 °C. Jurkat-Lucia NFAT-CD16 cells (Invivogen) (2 x 10^5^ cells/well and 1 x 10^5^ cells/well for spike and other protein respectively) were added and incubated for 24 hours at 37 °C, 5% CO2. Twenty mL of supernatant was then transferred to a white 96-well plate with 50 mL of reconstituted QUANTI-Luc secreted luciferase and read immediately on a Victor 3 luminometer with 1s integration time. Relative light units (RLU) of a no antibody control was subtracted as background. Palivizumab was used as a negative control, while CR3022 was used as a positive control, and P2B-2F6 to differentiate the Beta from the D614G variant. To induce the transgene 13 cell stimulation cocktail (Thermofisher Scientific, Oslo, Norway) and 2 µg/mL ionomycin in R10 was added as a positive control to confirm sufficient expression of the Fc receptor. CR3022 (for spike and RBD) or 4A8 (NTD) were used as positive controls and Palivizumab were used as negative controls. RLUs for original and Beta spikes were normalised to each other and between runs using CR3022. A cut off of 40 was determined by screening of 40 SARS-CoV-2 naive and unvaccinated individuals. All samples were run head to head in the same experiment as were all variants tested.

### Antibody-dependent cellular trogocytosis (ADCT) assay

ADCT was performed as described in and modified from a previously described study (Richardson, Crowther et al. 2018). HEK293T cells transfected with a SARS-CoV-2 spike pcDNA vector as above were surface biotinylated with EZ-Link Sulfo-NHS-LC-Biotin as recommended by the manufacturer. Fifty-thousand cells per well were incubated with 5-fold titration of mAb starting at 25 µg/mL or single 1 in 100 dilution for 30 minutes. Following an RPMI media wash, these were then incubated with carboxyfluorescein succinimidyl ester (CFSE) stained THP-1 cells (5 x 10^4^ cells per well) for 1 hour and washed with 15mM EDTA/PBS followed by PBS. Cells were then stained for biotin using Streptavidin-PE and read on a FACSAria II. Trogocytosis score was determined as the proportion of CFSE positive THP-1 cells also positive for streptavidin-PE less the no antibody control with waves run head-to-head.

### Antibody-dependent monocyte phagocytosis (ADMP) assay

Human monocytes (effector cells), purified from PBMCs using CD14 microbeads (Miltenyi), were employed in ADMP assays. Vero E6 cells were infected with SARS-CoV-2-mNG at an MOI of 0.1 for 48 h. On day 2, target cells, harvested with trypsin-EDTA (0.25%), were pre-incubated with tested antibodies at 37°C for 90 min. Monocytes were added to the target cells at a 4:1 ratio (effector:target) and cocultured for 4 h. The cells were then stained with CD14-PE (clone 63D3, BioLegend) and CD66b-APC (Clone G10F5, BioLegend) for 10 min, washed once with PBS, and fixed with 4% paraformaldehyde twice, following the BSL3 protocol. Cell acquisition was performed in LSR Fortessa, and FlowJo software version 10.8 (Tree Star) was used for analysis. Monocytes were identified as CD14+CD66b-SSC-Aint. The percent phagocytosis by monocytes was calculated as the frequency of mNG+ cells. A fold change in percent phagocytosis relative to prior infection was used to quantify ADMP induction in this study.

### SARS-CoV-2 VSV-G virus production

The generation of a replication-competent VSV expressing SARS-CoV-2 S protein with a 21 amino-acid C-terminal deletion that replaces the VSV G protein (VSV-SARS-CoV-2) was described previously (Case, Rothlauf et al. 2020). The S protein-expressing VSV virus was propagated in MA104 cell culture monolayers (African green monkey, ATCC CRL-2378.1) as described previously (Case, Rothlauf et al. 2020), and viral stocks were titrated on Vero E6 cell monolayer cultures. VSV plaques were visualized using neutral red staining. All work with infectious SARS-CoV-2 was performed in Institutional Biosafety Committee approved BSL3 and A-BSL3 facilities at Washington University School of Medicine using appropriate positive pressure air respirators and protective equipment.

### Animal studies

Animals were group housed in micro-isolator cages at beginning of study. Food and bedding were routinely supplied, changed, and monitored. Drinking water was provided ad libitum. Animals were acclimated to study housing for at least 72 hours prior to initiation of the studies.

#### Prophylactic studies

48–54-week-old female K18-hACE2 transgenic mice expressing the human ACE2 receptor were purchased from Jackson Laboratories. Eight mice per group were prophylactically administered through the intraperitoneal route with 12 mg/kg of the 54043-5 antibodies (wt or LALA-PG version) or with an isotype control antibody (#1664) or 5 mg/kg of a SARS-CoV-2 neutralizing control antibody (S309) or the vehicle only (PBS) and 24 hours later intranasally infected with 2×10^3^ PFU of SARS-CoV-2 (WA1/2020 strain). Weight and survival were monitored for 14 days. Three mice per group were sacrificed at day 3 p.i. and lungs collected for determining viral titers.

#### Therapeutic studies

9–10-week-old female BALB/cAnNHsd mice were purchased from Envigo. Eight mice per group were intranasally challenged with 10^5^ TCID50 of SARS-CoV-2 (MA10 strain) and 24 hours p.i. administered through the intraperitoneal route with 100 mg/kg of the 54043-5 antibodies (wt or LALA-PG version) or with 200 mg/kg of an isotype control (DENV-2D22) or 10 mg/kg of a SARS-CoV-2 human neutralizing antibody (COV2-2381). Weight, clinical signs, and survival were monitored for 5 days. Four mice per group were sacrificed at day 2 and 5 p.i. and lungs collected for determining viral titers, cytokine analysis and histopathology.

#### Lung viral titer determination

Upon collection, lung tissues were homogenized using a QIAGEN TissueLyser in 1 mL of DMEM supplemented with penicillin and streptomycin and clarified via centrifugation. Clarified supernatant was stored frozen (≤ -65°C) after removal of the aliquot for RNA extraction which was added to TRIzol® LS Reagent and mixed thoroughly for viral inactivation and stored frozen (≤ -65°C).

For qRT-PCR analysis, samples were removed from frozen storage, thawed, and processed for RNA extraction and purification using the Zymo Direct-zol^TM^ RNA Mini Prep kit. RNA samples were analyzed via quantitative RT-PCR using a Bio-Rad CFX96TM Real-Time PCR Detection System. Results are reported as viral genomic equivalent per milliliter (GEq/mL).

For TCID50 assay analysis, samples were removed from frozen storage and allowed to thaw under ambient conditions. Tissue homogenate samples were serially, 10-fold diluted in MEM/2% HI-FBS and used to infect Vero E6 cells in 96-well plates. These plates were cultured for 3 days before assessing viral titers based on microscopic cytopathic effects. Viral load data are expressed as TCID50 per gram of tissue.

#### Cytokine Analysis

For cytokine analysis, clarified supernatant from tissue homogenates were processed using the Bio-Plex Pro Mouse Cytokine 23-plex Assay kit (Bio-Rad).

#### Histopathology

Histopathologic analysis of lung tissues collected at day 2 and day 5 p.i. on the antibody therapeutic mouse studies was performed by the Experimental Pathology Laboratories Inc. (EPL) of the University of Texas. In brief, upon collection, lung tissues were perfused with formalin and placed in 10% neutral buffered formalin for fixation. Fixed tissues were shipped to EPL for embedding and hematoxylin and eosin processing along with completed copies of relevant necropsy documentation.

Mixed or mononuclear cell alveolar, bronchoalveolar and/or interstitial inflammation, mononuclear cell perivascular inflammation, alveolar hyperplasia, mesothelial hypertrophy, hemorrhage, and necrosis of bronchial and bronchiolar epithelium were considered related to SARS-CoV-2 infection.

## QUANTIFICATION AND STATISTICAL ANALYSIS

The ELISA error bars (± standard error of the mean) were calculated using Graphpad Prism 9.5.0. Mean ± SEM or mean ± SD were determined for continuous variables as noted. Technical and biological replicates are described in the relevant figure legends. Details of the statistical analysis can be found in the main text and respective figure captions.

For mouse studies, significance in the body weight differences was evaluated using an ordinary two-way ANOVA. A Dunnett’s multiple comparisons test with a single pooled variance computed for each comparison was utilized.

For lung viral titer analysis, significance of the differences in their levels was evaluated using an RM one-way ANOVA. Tukey’s multiple comparisons test with individual variances computed for each comparison was utilized.

For cytokine and chemokine analysis, significance of the differences in their levels was evaluated using an ordinary two-way ANOVA. Tukey’s multiple comparisons test with individual variances computed for each comparison was utilized.

For mouse studies, all statistical analysis was performed using GraphPad Prism V.9.00 software (San Diego, CA), and a *p* value <0.05 was considered statistically significant.

