## Supplemental Figures and Tables for "Discovery and Characterization of a Pan-betacoronavirus S2-binding antibody"

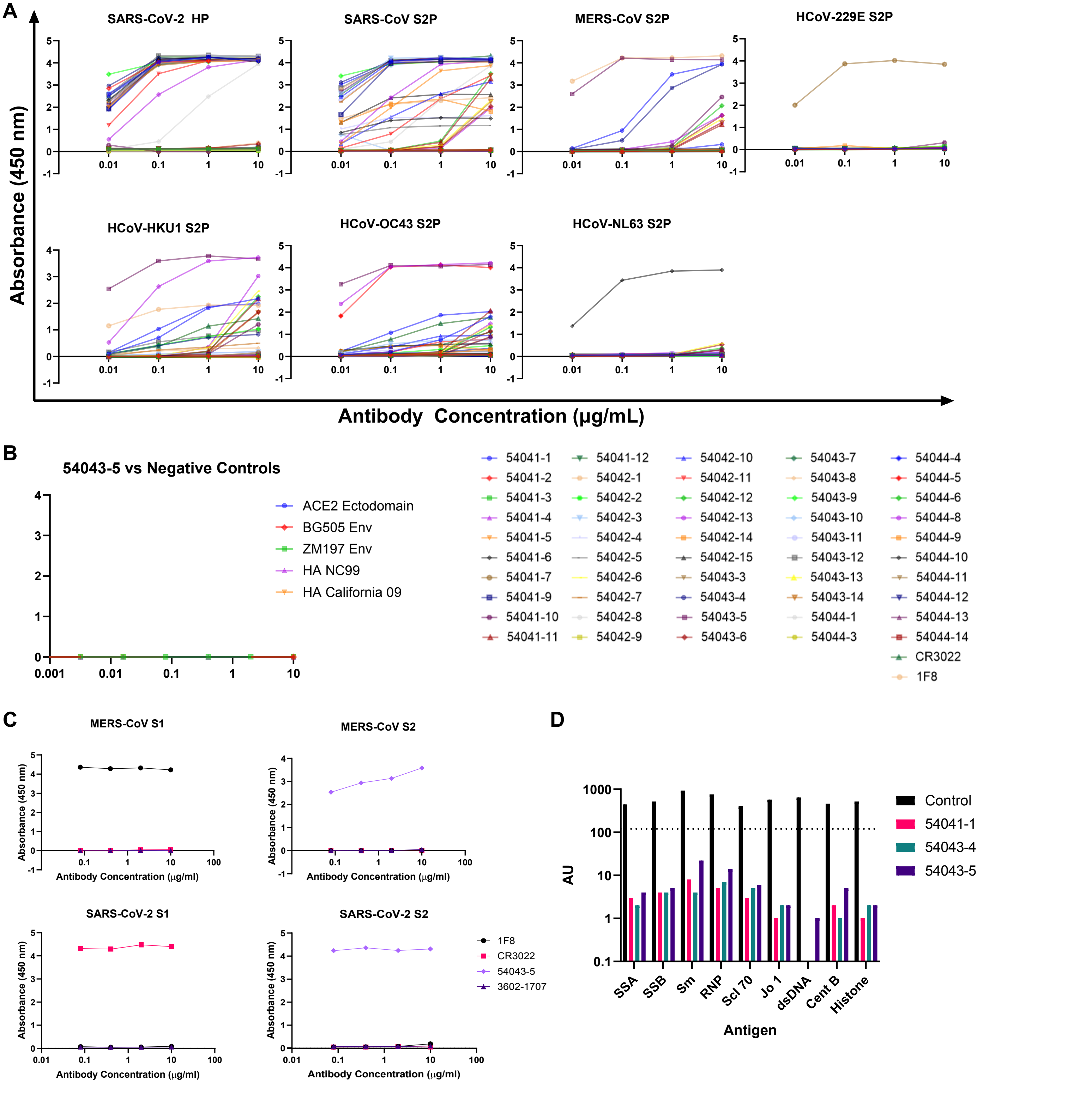


**Figure S1. ELISA Screening curves and 54043-5 SPR binding kinetics**
**(A)** Antibodies were tested for binding to a panel of CoV antigens by ELISA. The SARS-CoV-2/SARS-CoV cross reactive antibody CR3022 was used as a positive control for SARS-CoV-2 and SARS-CoV. The anti-MERS RBD antibody 1F8 was used as a positive control for MERS-CoV. All ELISAs were performed in technical duplicates with at least 2 biological duplicates. Data are represented as means ±SEMs. **(B)** 54043-5 was tested against a panel of non-coronavirus antigens to assess polyreactivity. **(C)** 54043-5 was screened in an ELISA against recombinant S1 and S2 subdomains of MERS-CoV and SARS-CoV-2. Anti-MERS antibody 1F8 was used as a positive control for MERS-CoV S1, and anti-SARS-CoV/SARS-CoV-2 antibody CR3022 was used as a positive control for SARS-CoV-2 S1. Anti-influenza antibody 3602-1707 served as a negative control. **(D)** mAb 54043-5 and two other broadly reactive lead mAbs identified in figure 1A were tested for autoreactivity against a panel of antigens in the Luminex AtheNA assay. AU stands for Athena units. Anti-HIV antibody 4E10 was used as a positive control for autoreactivity.

### Figure S2: Cryo-EM Data Processing

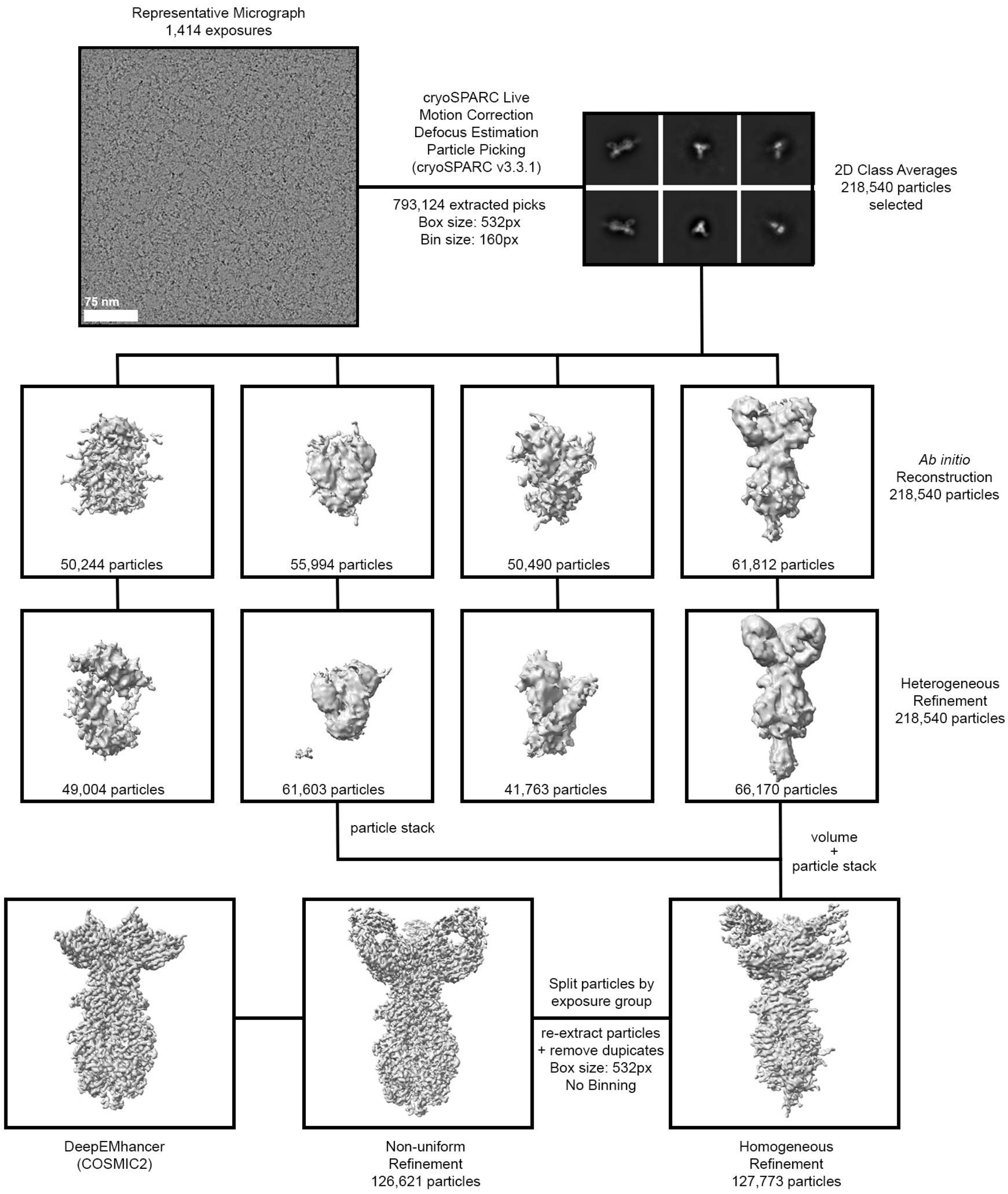

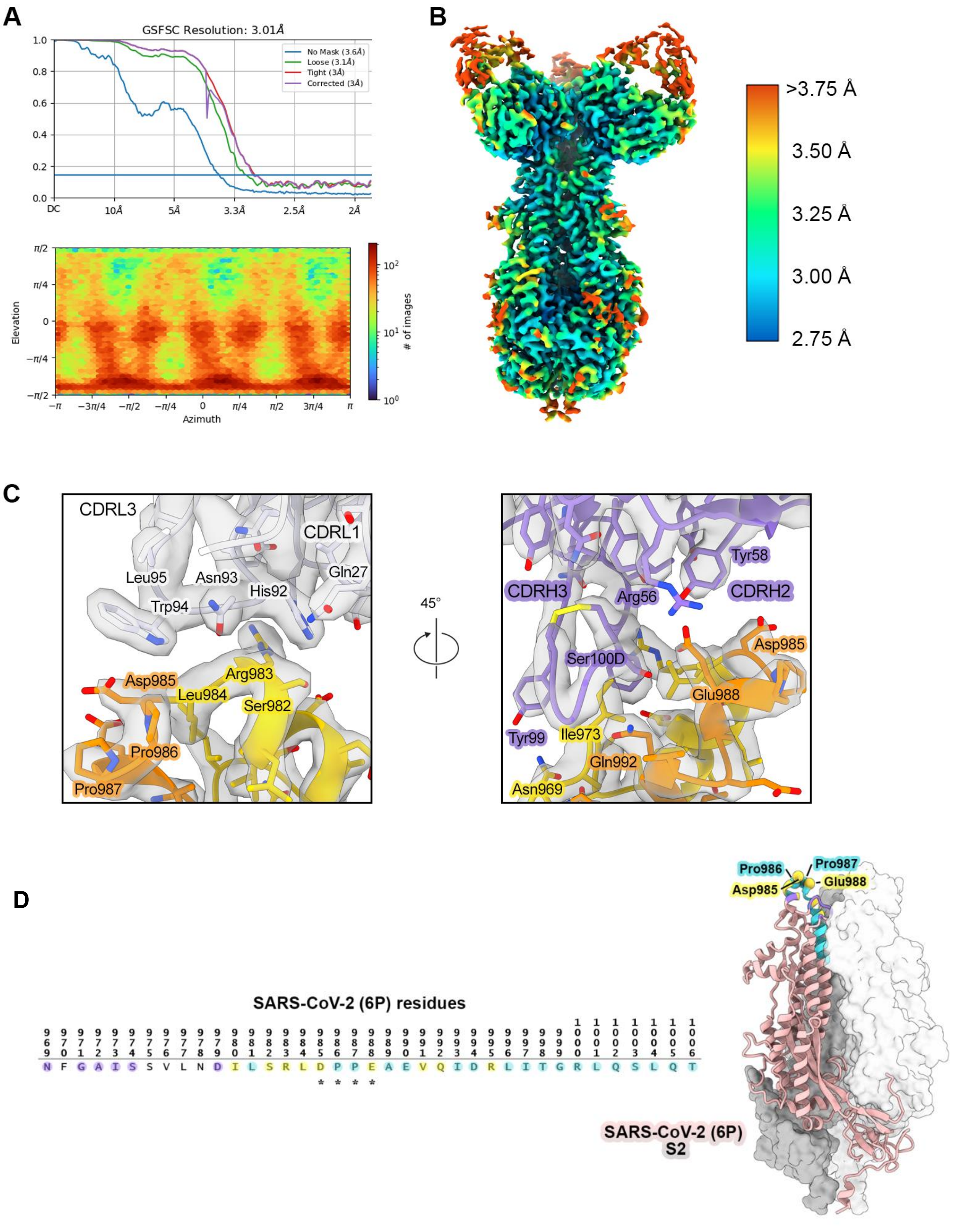


**Figure S3. Cryo-EM data validation and epitope comparison of 54043-5 and antibody 3A3 (A)** Gold-standard FSC curves (top) and particle orientation distribution (bottom) for the final refinement volume of Fab 54043-5 bound to SARS-CoV-2 S2. **(B)** Cryo-EM structure of Fab 54043-5 bound to SARS-CoV-2 S2, colored according to local resolution (Å). **(C)** Close-up views of the 54043-5 epitope, shown as cartoons colored according to Figure 3, with the corresponding map volume overlaid in partially transparent gray. Important interface residues are shown as sticks. Oxygen atoms are colored red, nitrogen atoms are colored blue, and sulfur atoms are colored yellow. **(D)** Left: Comparison of the 54043-5 epitope and residues identified to be part of the epitope for antibody 3A3 by hydrogen-deuterium exchange mass spectrometry and low-resolution cryo-EM (Silva et al., 2022). Residues within the 54043-5 epitope are highlighted in purple, residues within the 3A3 epitope are highlighted in cyan, and residues within both epitopes are highlighted in yellow. An "*" denotes residues that affected 3A3 binding to SARS-CoV-2 S when mutated (Silva et al., 2022). Right: The SARS-CoV-2 S2 subunit, with two protomers shown as gray surface and one protomer as pink cartoon. Residues at the apex of one protomer are colored as in (left) with * denoted residues labeled and Cα atoms shown as spheres.

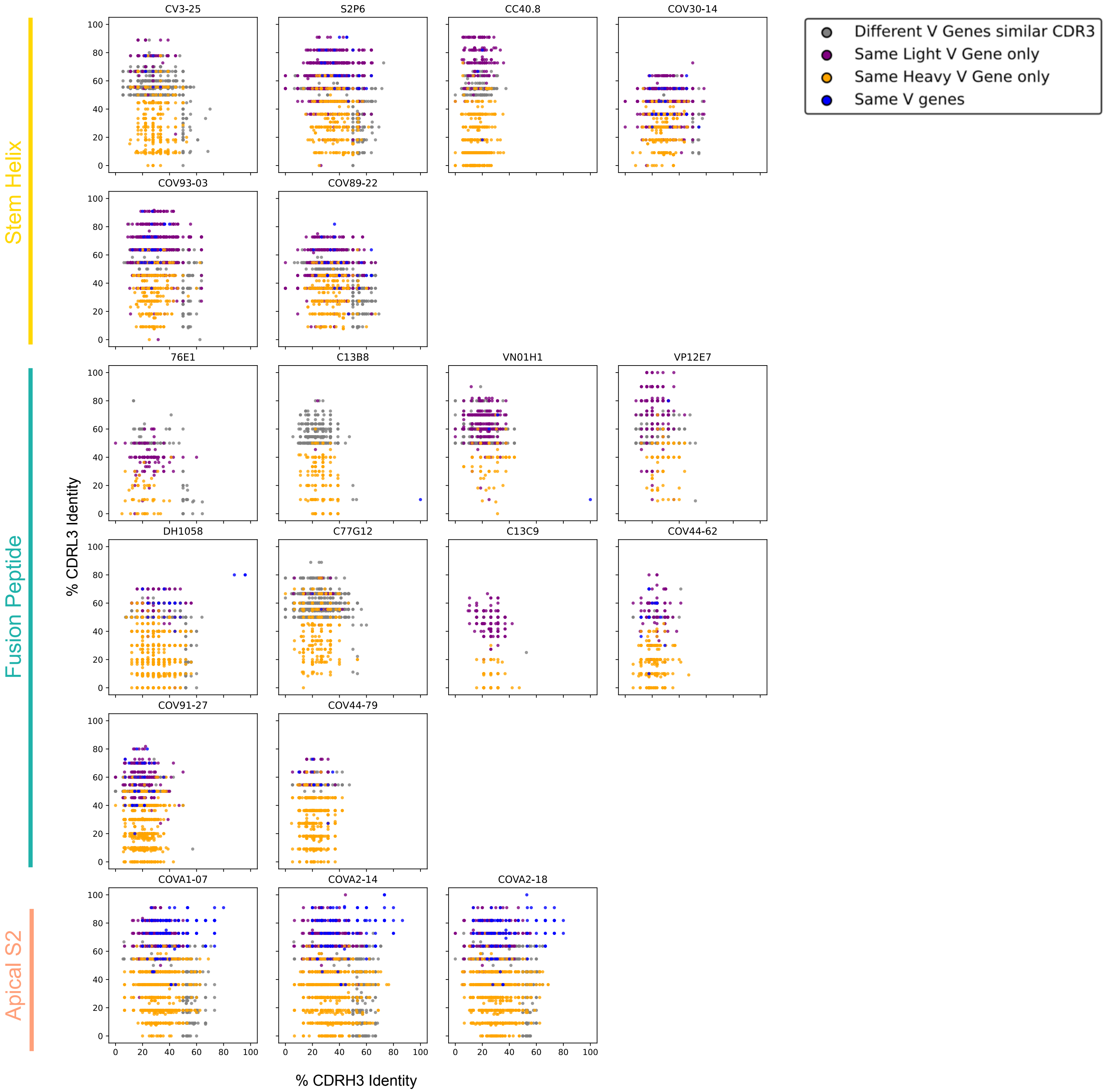


**Figure S4. Clonotype frequency of S2-binding antibodies**
The plots compare all antibodies from the CoVAbDab (each as a dot) against a reference S2-directed antibody, indicated above each plot. The dot's position reflects the CDRH3 (x-axis) and CDRL3 (y-axis) amino acid identity to the S2 antibody, with color coding indicating shared V gene usage. Inclusion criteria for dots are at least one matching V gene or ≥50% CDR3 sequence identity with the reference S2 antibody.

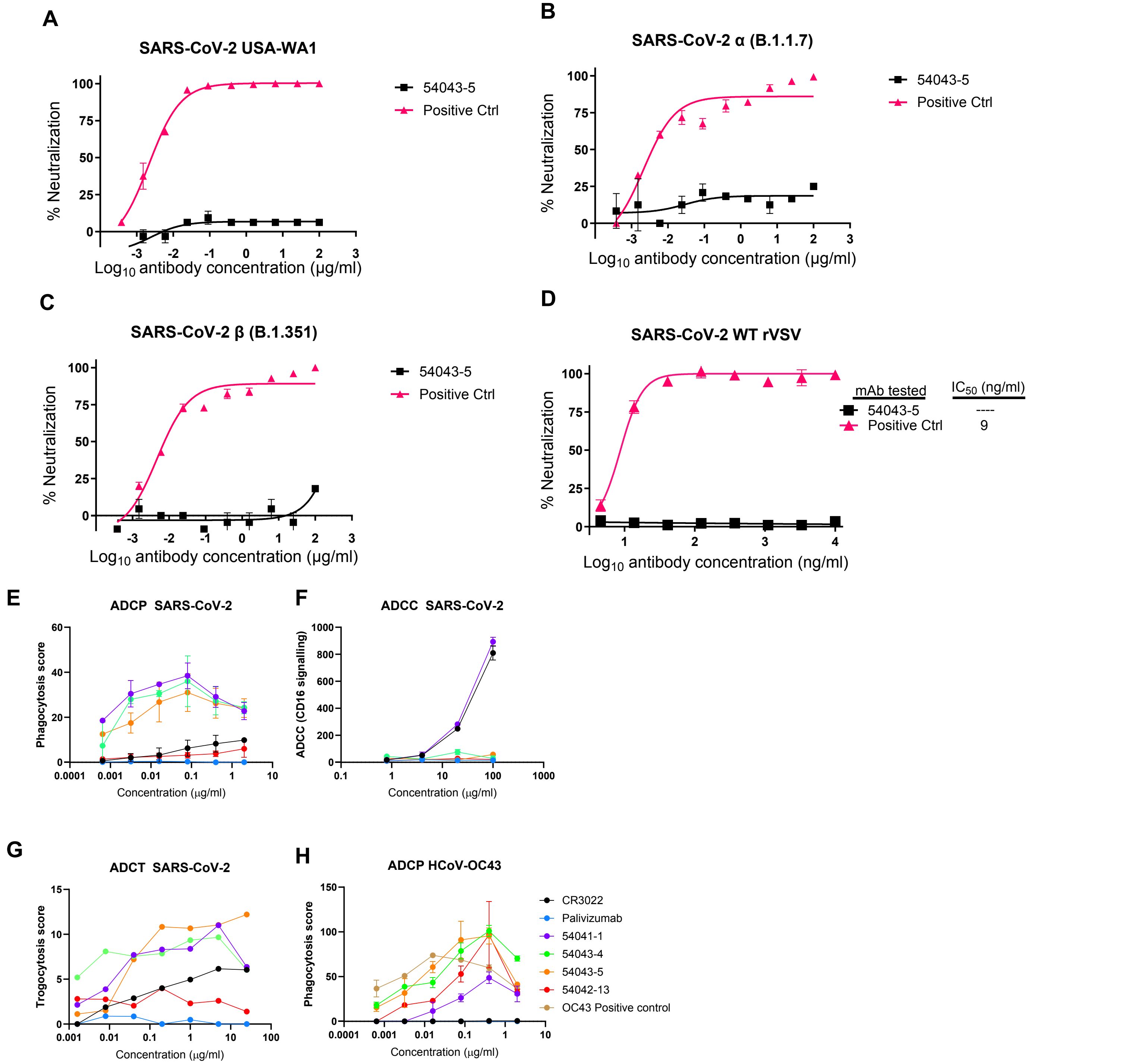


**Figure S5. Live virus and VSV-G pseudovirus neutralization experiments and Fc effector function data**
**(A-C)** Live virus neutralization curves for 54043-5 against SARS-CoV-2 USA WA-1, SARS-CoV-2 Alpha, SARS-CoV-2 Beta. **(D)** SARS-CoV-2 VSV-G neutralization curves for mAb 54043-5. Positive control is mAb 54042-4. The IC50 values were calculated in GraphPad Prism software by four-parameter best-fit. Data represent the percentage of neutralization as the mean ± SD; data are representative of at least two independent experiments performed in technical duplicate **(E)** Lead cross-reactive mAb candidates, 54041-1, 54043-4, 54043-5, and 54042-13 were tested for their ability to mediate antibody-dependent cellular phagocytosis (ADCP) for SARS-CoV-2, compared to antigen-positive control CR3022 and negative control palivizumab (an anti-RSV antibody). Phagocytosis score (methods) is shown on the y-axis and antibody concentration is shown on the x-axis. **(F)** Cross-reactive mAbs were tested for their ability to mediate antibody-dependent cellular cytotoxicity (ADCC) against SARS-CoV-2, compared to antigen-positive control CR3022 and negative control palivizumab. ADCC/CD16 signaling (methods) is shown on the y-axis and antibody concentration is shown on the x-axis. **(G)** Cross-reactive mAbs were tested for their ability to mediate antibody-dependent cellular trogocytosis (ADCT) against SARS-CoV-2, compared to antigen-positive control CR3022 and negative control palivizumab. Trogocytosis score (methods) is shown on the y axis and mAb concentration is shown on the x-axis **(H)** Cross-reactive mAbs were additionally tested against OC43 for their ability to mediate antibody-dependent cellular phagocytosis compared to the OC43-specific positive control, 54044-5, and negative control palivizumab. All Fc effector data is shown as mean ±SDs.

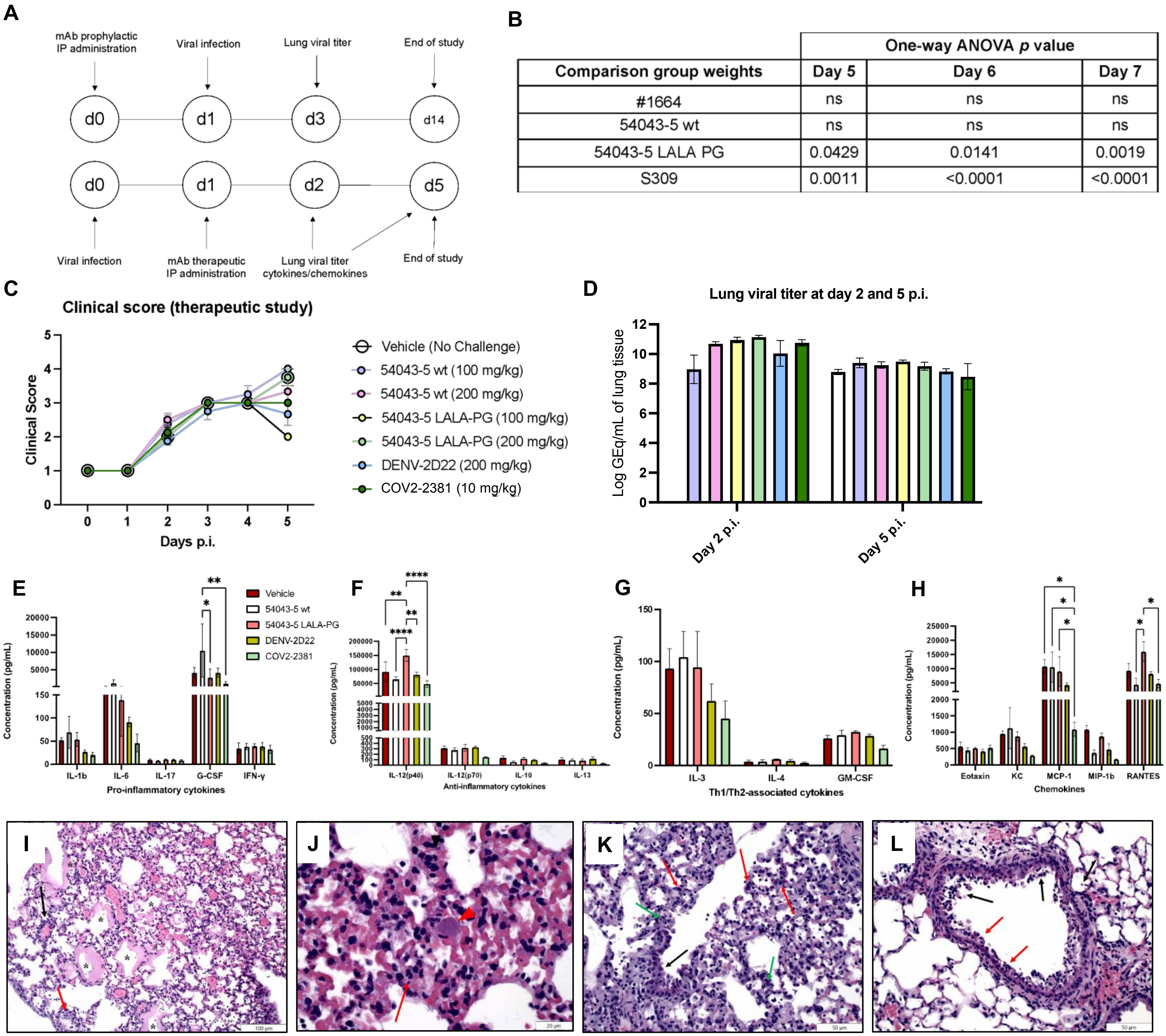


**Figure S6. *In vivo* efficacy of mAb 54043-5 in mouse models**

**(A)** Experimental workflow of the prophylactic and therapeutic challenge studies in Figure 6. **(B)** Values for statistical significance of the body weight data for mice in the prophylactic study, related to Figure 6A. **(C)** Clinical score values for mice in the therapeutic study, monitored for days 0-5 post-infection (p.i.). Results are expressed as absolute mean values plus SEM. **(D)** Lung viral titers of mice sacrificed on day 2 and 5 p.i. expressed as the logarithm of the number of viral genomic equivalents (GEq) per mL of homogenized lung tissue. Results are expressed as absolute mean values plus SEM. **(E-H)** Pro-inflammatory **(E)**, anti-inflammatory **(F)**, Th1/Th2-associated **(G)** cytokines and chemokines **(H)** levels from homogenized lung tissues collected from mice in the therapeutic study at day 5 p.i. Results are expressed as absolute mean values plus SEM. ****, p<0.0001; ***, p<0.001; **, p<0.01; *, p<0.05. **(I-L)** Representative histopathological examinations of lung tissues from deceased SARS-CoV-2 challenged mice in the therapeutic study. **(I)** Mild (grade 2) bronchointerstitial pneumonia and mild (grade 2) vasculitis. Mixed inflammatory cells expand alveolar septa (red arrow) and extend into alveolar airways. Multiple alveolar airways are enlarged and lined by thick hyaline membranes (*). Low numbers of inflammatory cells are present within the wall and along the endothelium of a small vessel (black arrow). **(J)** 5X magnification of panel A. Alveolar septa are expanded by mixed inflammatory cells (red arrow), and a fibrin microthrombus is present within an alveolar capillary (red arrowhead). **(K)** Moderate (grade 3) bronchointerstitial pneumonia, mild (grade 2) type II pneumocyte proliferation and mild (grade 2) bronchiolar hyperplasia. The alveolar septa are lined by numerous hyperplastic type II pneumocytes (red arrows), and alveolar septa are expanded by mixed inflammatory cells (green arrows). Bronchiolar hyperplasia is indicated by focal crowding of enlarged epithelial cells lining a terminal bronchiole (black arrow). **(L)** Mild (grade 2) bronchiolar necrosis. Necrotic epithelial cells (red arrows) are present around the circumference of a bronchiole and there is exfoliation of degenerating/necrotic epithelial cells (black arrows).

Table S1: Heatmap of amino acid percent identies of pairs of coronavirus spike proteins used in this study. The table is shaded from white being 100% amino acid identity to dark red being the lowest percent identity in the table.

| | SARS2 WA1 | SARS2 D614G | SARS2 Alpha | SARS2 Beta | SARS2 Delta | SARS2 Omicron BA.1 | SARS2 Omicron BA.2 | WIV1 | SARS1 | HKU9 | OC43 | MERS | HKU1 | 229E | SDCV | NL63 | PEDV | TGEV |
|---|---|---|---|---|---|---|---|---|---|---|---|---|---|---|---|---|---|---|
| **SARS2 WA1** | 100 | 99.92 | 99.41 | 99.33 | 99.5 | 97.23 | 97.65 | 77.96 | 76.75 | 34.01 | 33.01 | 32.03 | 31.67 | 27.9 | 27.37 | 27.11 | 26.67 | 26.16 |
| **SARS2 D614G** | 99.92 | 100 | 99.5 | 99.41 | 99.58 | 97.31 | 97.73 | 77.87 | 76.67 | 34.1 | 33.01 | 32.03 | 31.67 | 27.9 | 27.37 | 27.11 | 26.67 | 26.16 |
| **SARS2 Alpha** | 99.41 | 99.5 | 100 | 99.07 | 99.16 | 97.14 | 97.56 | 77.6 | 76.46 | 33.92 | 32.97 | 31.86 | 31.49 | 28.02 | 27.5 | 27.19 | 26.56 | 26.05 |
| **SARS2 Beta** | 99.33 | 99.41 | 99.07 | 100 | 98.99 | 97.14 | 97.56 | 77.77 | 76.56 | 34.1 | 33 | 32.06 | 31.79 | 28.18 | 27.33 | 27.19 | 26.75 | 26.14 |
| **SARS2 Delta** | 99.5 | 99.58 | 99.16 | 98.99 | 100 | 97.14 | 97.65 | 78.01 | 76.8 | 34.16 | 33.24 | 31.74 | 31.81 | 27.85 | 27.37 | 27.16 | 26.72 | 26.02 |
| **SARS2 Omicron BA.1** | 97.23 | 97.31 | 97.14 | 97.14 | 97.14 | 100 | 98.57 | 76.61 | 75.58 | 33.81 | 32.91 | 31.51 | 31.34 | 27.49 | 27.25 | 26.77 | 26.61 | 25.82 |
| **SARS2 Omicron BA.2** | 97.65 | 97.73 | 97.56 | 97.56 | 97.65 | 98.57 | 100 | 76.55 | 75.77 | 33.66 | 32.91 | 31.68 | 31.48 | 27.88 | 27.16 | 27 | 26.47 | 25.77 |
| **WIV1** | 77.96 | 77.87 | 77.6 | 77.77 | 78.01 | 76.61 | 76.55 | 100 | 92.53 | 33.6 | 32.76 | 31.35 | 31.75 | 27.53 | 26.18 | 25.21 | 25.91 | 26.3 |
| **SARS1** | 76.75 | 76.67 | 76.46 | 76.56 | 76.8 | 75.58 | 75.77 | 92.53 | 100 | 33.51 | 32.76 | 31.69 | 31.8 | 27.71 | 25.18 | 25.69 | 25.34 | 26.3 |
| **HKU9** | 34.01 | 34.1 | 33.92 | 34.1 | 34.16 | 33.81 | 33.66 | 33.6 | 33.51 | 100 | 30.34 | 29.61 | 30.5 | 25.05 | 25.3 | 23.5 | 23.26 | 23.11 |
| **OC43** | 33.01 | 33.01 | 32.97 | 33 | 33.24 | 32.91 | 32.91 | 32.76 | 32.76 | 30.34 | 100 | 31.25 | 63.42 | 25.71 | 26.13 | 24.68 | 24.8 | 24.93 |
| **MERS** | 32.03 | 32.03 | 31.86 | 32.06 | 31.74 | 31.51 | 31.68 | 31.35 | 31.69 | 29.61 | 31.25 | 100 | 29.95 | 26.01 | 24.06 | 23.7 | 24.35 | 23.49 |
| **HKU1** | 31.67 | 31.67 | 31.49 | 31.79 | 31.81 | 31.34 | 31.48 | 31.75 | 31.8 | 30.5 | 63.42 | 29.95 | 100 | 26.8 | 25.15 | 25.17 | 24.63 | 25.55 |
| **229E** | 27.9 | 27.9 | 28.02 | 28.18 | 27.85 | 27.49 | 27.88 | 27.53 | 27.71 | 25.05 | 25.71 | 26.01 | 26.8 | 100 | 44.03 | 64 | 49.67 | 48.83 |
| **SDCV** | 27.37 | 27.37 | 27.5 | 27.33 | 27.37 | 27.25 | 27.16 | 26.18 | 25.18 | 25.3 | 26.13 | 24.06 | 25.15 | 44.03 | 100 | 43.49 | 44.73 | 42.07 |
| **NL63** | 27.11 | 27.11 | 27.19 | 27.19 | 27.16 | 26.77 | 27 | 25.21 | 25.69 | 23.5 | 24.68 | 23.7 | 25.17 | 64 | 43.49 | 100 | 44.3 | 46.77 |
| **PEDV** | 26.67 | 26.67 | 26.56 | 26.75 | 26.72 | 26.61 | 26.47 | 25.91 | 25.34 | 23.26 | 24.8 | 24.35 | 24.63 | 49.67 | 44.73 | 44.3 | 100 | 46.61 |
| **TGEV** | 26.16 | 26.16 | 26.05 | 26.14 | 26.02 | 25.82 | 25.77 | 26.3 | 26.3 | 23.11 | 24.93 | 23.49 | 25.55 | 48.83 | 42.07 | 46.77 | 46.61 | 100 |

Supplemental Table S2

#### EM data collection

| | |
|---|---|
| Microscope | FEI Talos Glacios |
| Voltage (kV) | 200 |
| Detector | Falcon 4 |
| Magnification (nominal) | 150,000 |
| Pixel size (Å/pix) | 0.94 |
| Exposure rate ($e^-$/pix/sec) | 6 |
| Frames per exposure | 60 |
| Exposure ($e^-$/Å$^2$) | 50 |
| Defocus range (μm) | 1.5-2.5 |
| Tilt angle (˚) | 0 |
| Micrographs collected | 1,885 |
| Micrographs used | 1,414 |
| Particles extracted (total) | 793,124 |
| Automation software | SerialEM |
| Sample | SARS-CoV-2 S2-37 + 54043-5 Fab |

#### 3D reconstruction statistics

| | |
|---|---|
| Particles | 126,621 |
| Symmetry | C3 |
| Map sharpening B-factor | -83 |
| Unmasked resolution at 0.5 FSC (Å) | 3.4 |
| Masked resolution at 0.5 FSC (Å) | 3.3 |
| Unmasked resolution at 0.143 FSC (Å) | 3.0 |
| Masked resolution at 0.143 FSC (Å) | 3.0 |

#### Model refinement and validation statistics

| | |
|---|---|
| Refinement package | Phenix |
| Refinement tool | Real-space refinement |
| Refinement strategies | min global, local_grid_search, adp, ss restraints, rotamer restraints, Ramachandran restraints |
| **Composition** | |
| Amino acids | 2040 |
| RMSD bonds (Å) | 0.005 |
| RMSD angles (°) | 0.645 |
| **Average B-factors** | |
| Amino acids | 57.45 |
| **Ramachandran** | |
| Favored (%) | 97.08 |
| Allowed (%) | 2.92 |
| Outliers (%) | 0 |
| Rotamer outliers (%) | 0 |
| Clash score | 3.09 |
| C-beta outliers (%) | 0 |
| CaBLAM outliers (%) | 1.50 |
| CC (mask) | 0.80 |
| MolProbity score | 1.26 |
| EMRinger score | 4.12 |

**Supplemental Table 2: Cryo-EM data collection, processing, and PDB model validation, related to Figure 3.**

Table S3: Epitope and paratope buried surface areas in the interaction of 54043-5 with SARS-CoV-2 Spike.
**(A)** The average amount of solvent accessible surface area in Å$^2$ buried by 54043-5 on each epitope residue of SARS-CoV-2 S over the three protomers in the structure.

**(B)** The average amount of solvent accessible surface area in Å$^2$ buried on 54043-5 by SARS-CoV-2 S on each antibody paratope residue over the three antibodies in the structure.

A

54043-5 epitope on SARS-CoV-2 S

| S Residue # | AA | Buried Surface Area (Å$^2$) |
|---|---|---|
| 755 | Gln | 16 |
| 969 | Asn | 18 |
| 971 | Gly | 10 |
| 972 | Ala | 10 |
| 973 | Ile | 97 |
| 974 | Ser | 10 |
| 979 | Asp | 23 |
| 980 | Ile | 2 |
| 982 | Ser | 57 |
| 983 | Arg | 173 |
| 984 | Leu | 33 |
| 985 | Asp | 61 |
| 986 | Pro | 8 |
| 988 | Glu | 96 |
| 991 | Val | 9 |
| 992 | Gln | 42 |
| 995 | Arg | 7 |

B

54043-5 paratope on SARS-CoV-2 S

| | Ab residue # | AA | Buried Surface Area (Å$^2$) |
|---|---|---|---|
| **Heavy Chain** | 52 | Tyr | 1 |
| | 56 | Arg | 13 |
| | 58 | Tyr | 38 |
| | 99 | Tyr | 76 |
| | 100A | Ala | 29 |
| | 100B | Gly | 40 |
| | 100D | Ser | 72 |
| | 100E | Cys | 13 |
| | 100F | Phe | 128 |
| | 100G | Leu | 5 |
| | 100H | Asn | 1 |
| **Light Chain** | 27 | Gln | 17 |
| | 30 | Ser | 9 |
| | 32 | Asn | 24 |
| | 91 | Tyr | 1 |
| | 92 | His | 70 |
| | 93 | Asn | 31 |
| | 94 | Trp | 69 |
